# Tunable Macroscopic Alignment of Self-Assembling Peptide Nanofibers

**DOI:** 10.1101/2024.02.02.578651

**Authors:** Adam C. Farsheed, Christian Zevallos-Delgado, Le Tracy Yu, Sajede Saeidifard, Joseph W.R. Swain, Jonathan T. Makhoul, Adam J. Thomas, Carson C. Cole, Eric Garcia Huitron, K. Jane Grande-Allen, Manmohan Singh, Kirill V. Larin, Jeffrey D. Hartgerink

## Abstract

Fibrous proteins that comprise the extracellular matrix (ECM) guide cellular growth and tissue organization. A lack of synthetic strategies able to generate aligned, ECM-mimetic biomaterials has hampered bottom-up tissue engineering of anisotropic tissues and led to a limited understanding of cell-matrix interactions. Here, we present a facile extrusion-based fabrication method to produce anisotropic, nanofibrous hydrogels using self-assembling peptides. The application of shear force coinciding with ion-triggered gelation is used to kinetically trap supramolecular nanofibers into aligned, hierarchical structures. We establish how modest changes in phosphate buffer concentration during peptide self-assembly can be used to tune their alignment and packing. In addition, increases in the nanostructural anisotropy of fabricated hydrogels are found to enhance their strength and stiffness under hydrated conditions. To demonstrate their utility as an ECM-mimetic biomaterial, aligned nanofibrous hydrogels are used to guide directional spreading of multiple cell types, but strikingly, increased matrix alignment is not always correlated with increased cellular alignment. Nanoscale observations reveal differences in cell-matrix interactions between variably aligned scaffolds and implicate the need for mechanical coupling for cells to understand nanofibrous alignment cues. In total, innovations in the supramolecular engineering of self-assembling peptides allow us to generate a gradient of anisotropic nanofibrous hydrogels, which are used to better understand directed cell growth.

## Introduction

Anisotropic, or directionally dependent, tissues contain aligned and hierarchical extracellular matrix (ECM) arrangements related to tissue function.^1,2^ Naturally, scientists and engineers have aimed to synthetically fabricate aligned fibrous scaffolds to direct cellular organization.^3,4^ Fiber spinning techniques such as electrospinning^5–7^ (with the recent addition of rotary jet spinning^8^) have primarily been used towards this goal, representing a bottom-up approach where individual fibers are deposited onto a rotating mandrel to form an aligned matrix. These methods, though, require specialized equipment, commonly involve the use of toxic solvents, and have difficulty recreating hierarchical 3D geometries. Alternatively, top-down extrusion-based techniques that exert shear forces onto flowing extrudates have successfully been used to align biopolymers such as collagen,^9–11^ but biologically-derived materials suffer from batch-to-batch variability, have sustainable sourcing concerns, and are difficult to chemically modify.^12^ Similar shear-based strategies involving synthetic materials have relied on two component systems, where preformed fibers have been embedded into hydrogels prior to extrusion.^13–15^ These materials, though, do not accurately recreate the fibrous nanostructure or chemical complexity of native ECM.

Self-assembling peptides, which rely on supramolecular mechanisms for assembly, represent a synthetical class of fibrous, ECM-mimetic biomaterials that have the benefits of design flexibility, biocompatibility, and chemical purity.^16,17^ A longstanding goal within the field of supramolecular chemistry has been to follow a biomimetic route starting at molecular design and resulting in macroscopic, hierarchical fibrous alignment.^18–20^ Extrusion of self-assembling peptide solutions into ionic gelation baths has allowed progress toward this goal, but the tunability of previously designed systems is severely limited.^21–26^ They have primarily been designed and characterized as having a binary switch between isotropy and anisotropy, and the inability to modulate the degree of alignment has hampered their optimization for biological applications.

Here, we synthetically fabricate aligned nanofibrous hydrogels with tunable alignments using a β-sheet-forming peptide called “K2” with sequence K_2_(SL)_6_K_2_(Figure 1a). Designed in our lab over a decade ago,^27^ K2 is part of the Multidomain Peptide family of assemblers.^28,29^ The amphiphilic repeat of K2 drives β-sheet formation via hydrophobic packing of the leucine residues reinforced by backbone hydrogen bonding, whereas charge-charge repulsion at each termini opposes assembly. The delicate balance of these competing interactions allows ionic strength to act as a self-assembly trigger. At low ionic strength conditions, a small number of K2 monomers dimerize, a small number of nanofibers form, and the solution remains a liquid. At high ionic strength conditions, charge screening of the lysine residues allows for widespread β-sheet nanofiber formation, and a bulk transition to a hydrogel (Figure 1a).

**Figure 1.**
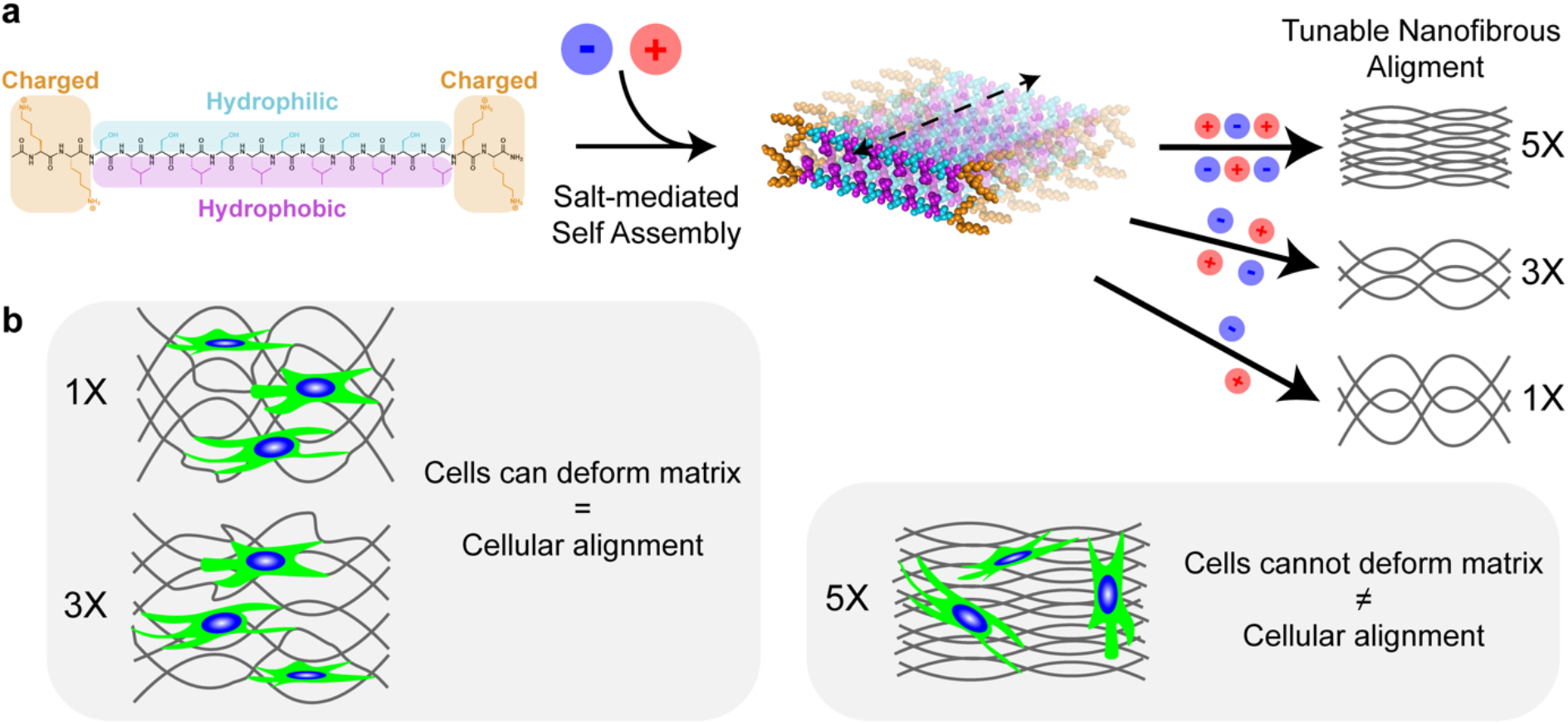
K2 Assembly and Cellular Spreading Schematic. (a) K2 structure, with charged (orange), hydrophilic (blue), and hydrophobic (purple) amino acids color coded. Salt-mediated β-sheet formation allows for tunable nanofibrous alignment. (b) Summary of cellular behavior on differentially aligned K2 matrices.

K2 forms a viscoelastic, shear-thinning hydrogel when dissolved in a physiological buffer, but counterintuitively, does not gain anisotropic properties when sheared.^30^ A mechanism explaining this phenomenon with self-assembling β-hairpin peptides has been investigated, in which fibrous domains fracture and recover under shear without affecting fibrillar alignment.^31^ Therefore, we decoupled the hydrogel formation process into two components: 1) A precursor solution containing K2 dissolved in Milli-Q water; 2) A gelation bath containing salts and a buffer component. The extrusion of the precursor solution into the gelation bath provides the necessary counterions for K2 nanofibrillar formation and the bulk transition from a liquid to a hydrogel (Movie S1). We imaged this process with polarized light microscopy (PLM) and observed birefringence within the resulting hydrogel, suggestive of structural anisotropy (Movie S2). In this work, we demonstrate how gelation bath composition determines K2 nanofibrous alignment, correlate hydrogel structural differences with bulk mechanical properties, and use this biomaterial system to better understand directed cellular spreading.

## Results and Discussion

### Investigation of Alignment Parameters

We initially sought to understand how the precursor solution’s K2 concentration influences the resulting hydrogel’s properties. Using a standard 100-1000 μL pipette tip and a phosphate-buffered saline (PBS) gelation bath, PLM revealed that no hydrogel formed using a 1wt% K2 precursor solution and that an unstable amorphous hydrogel with slight birefringence formed using a 2wt% K2 precursor solution (Figure S2a,b). In contrast, use of 3 or 4wt% K2 precursor solutions resulted in cylindrical, birefringent hydrogels (Figure S2c,d). These data suggest that K2 must be above a concentration threshold to form a robust, birefringent hydrogel, which is consistent with previously reported liquid crystalline behavior in other self-assembling peptide systems.^20^

While performing this study using a 3wt% K2 precursor solution, we observed an unexpected increase in viscosity over time when stored at 4°C (Figure S3a). Cryogenic transmission electron microscopy (Cryo-TEM) revealed that K2 nanofiber density increased after 1 day at 4°C, suggesting that the supramolecular equilibration in this system is on the order of days (Figure S3d). Further supporting these findings, attenuated total reflectance Fourier transform infrared spectroscopy (ATR-FTIR) showed an increase in the amide I band at 1616 cm^-1^, corresponding to C=O stretching within the β-sheet (Figure S3b). In addition, dynamic light scattering (DLS) showed a significant increase in particle size between day 0 and day 1 (Figure S3c). Further analysis of the 3wt% K2 precursor solution at day 21 revealed that an equilibrium had been reached by day 7 (Figure S3a-c). We therefore used 3wt% K2 that had been allowed to equilibrate at 4° C for >7 days as the precursor solution for the remainder of the study.

Next, we examined how the gelation bath composition affects the resulting hydrogel’s properties. Using a standard 0.1-10 μL pipette tip to screen a series of pH 7 gelation baths, PLM revealed a positive correlation between ionic strength and hydrogel birefringence (Figure S4). In addition, by manually lifting hydrogels from solution and observing if they fractured, a positive correlation between phosphate buffer concentration and hydrogel strength was found (Figure S4 checkmarks). Further implicating the importance of gelation bath composition, the precursor solution extrusion rate did not affect the observed birefringence pattern within the same gelation bath (Figure S5).

We were interested in further investigating differences between the 10 mM phosphate buffer/ 140 mM NaCl gelation bath, henceforth referred to as 1X PB, and the 50 mM phosphate buffer/ 140 mM NaCl gelation bath, henceforth referred to as 5X PB, due to them leading to differences in hydrogel birefringence, strength, and being close in composition to phosphate-buffered saline (PBS) commonly used in cell culture. We therefore added 3X PB (30 mM phosphate buffer/ 140 mM NaCl, pH 7) and 10X PB (100 mM phosphate buffer/ 140 mM NaCl, pH 7) gelation baths to analyze a spectrum of phosphate buffer concentrations. In addition, a “pregelled” 3wt% K2 unaligned control was included, in which 3wt% K2 was dissolved in Hank’s Balanced Salt Solution (HBSS) to form a hydrogel^30^ and then extruded into 1X PB. PLM revealed a positive correlation between PB concentration and hydrogel birefringence at multiple extrusion diameters, although minor differences between groups were difficult to discern (Figure S6). We therefore used scanning electron microscopy (SEM) and 2D small-angle x-ray scattering (2D SAXS) to carry out a more robust analysis of hydrogels extruded through a standard 100-1000 μL pipette tip (Figure 2). Regardless of alignment parameters, an expansive network of nanofibers was observed, while PB concentration was qualitatively found to positively correlate with an increase in fiber alignment and packing (Figure 2b,c). In addition, close observation revealed an increase in nanofiber bundling as a function of PB concentration (Figure 2c). 2D SAXS quantitatively confirmed the trends observed via PLM and SEM, as the full width at half maximum (FWHM) of the peak within the azimuthal distribution decreased from 71° to 61° to 53° between 1X, 3X, and 5X PB hydrogels, respectively (Figure 2d,e). Thus, we successfully engineered a self-assembling peptide system by which modest changes in phosphate buffer can be used to tune hierarchical nanofibrillar alignment (Figure 1a).

**Figure 2.**
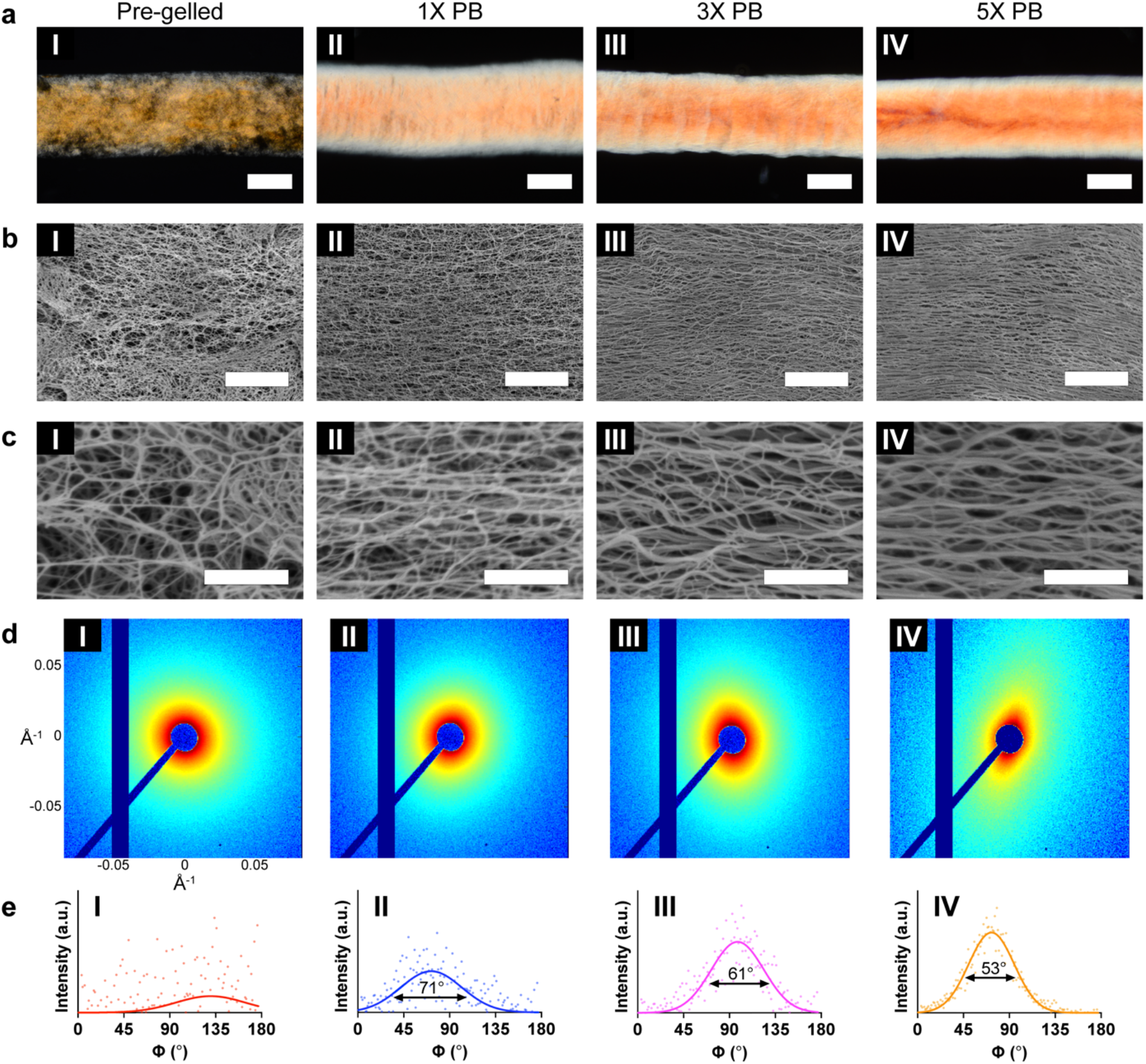
Tunable Alignment of Nanofibrous K2 Hydrogels. (a) Polarized light micrographs (scale bars = 500 μm), (b, c) scanning electron micrographs (scale bars = 3 μm and 500 nm, respectively), (d) 2D small-angle x-ray scattering patterns, and (e) azimuthal distributions derived from 2D small-angle x-ray scattering patterns of (I) pre-gelled, (II) 1X PB, (III) 3X PB, and (IV) 5X PB hydrogels. The full width at half maximum for the largest peak is indicated underneath the curve.

After seeing their effects on K2 nanofibrillar alignment, we were interested in understanding how gelation bath composition affects bulk hydrogel geometry. We expected that hydrogel diameters would match the inner diameters of the tips used for extrusion (860 and 430 μm for 100-1000 and 0.1-10 μL pipette tips, respectively). This was true for pre-gelled hydrogels (813 and 440 μm), but the 1X, 3X, and 5X PB hydrogels followed an unexpected trend (Figure S7): 1X PB hydrogels were significantly larger (1156 and 618 μm), 3X PB hydrogels were similar (780 and 363 μm), and 5X PB hydrogels were significantly smaller (717 and 333 μm) when compared to pre-gelled hydrogels.

These data help elucidate a diffusion-mediated mechanism underlying the positive correlation between PB concentration and K2 nanofibrillar alignment. Specifically, it helps in understanding what occurs between when the liquid K2 extrudate leaves the pipette tip and when ions within the gelation bath physically crosslink enough K2 nanofibers to initiate bulk gelation. We hypothesize that there are two competing diffusive interactions within the liquid extrudate during this time: **1**) the outward diffusion of high-concentration K2 monomers toward the gelation bath; **2**) the outward diffusion of water molecules toward the high-ionic strength gelation bath (dehydration). Macroscopically, **1** causes an increase in hydrogel diameter whereas **2** causes a decrease in hydrogel diameter. Thus, as PB concentration increases, the relative rate of **2** increases, which explains why hydrogels prepared in 1X (**1** > **2**), 3X (**1** = **2**), and 5X PB (**1** < **2**) gelation baths have declining diameters. In addition, this mechanism explains why the diameter of 5X PB hydrogels is smaller than the pipette tip inner diameter.

Considering why PB concentration leads to the observed differences in nanoscale geometry, the greater **2** compared to **1**, the higher nanofiber bundling and alignment is expected. This is in agreement with the mechanism for thermally-mediated peptide amphiphile bundling and alignment,^21^ which also implicates dehydration as the primary driver. Thus, although the shear forces present during extrusion are consistent regardless of PB concentration, higher alignment is achieved for higher PB concentrations. By pairing macroscopic and nanoscale observations, we can propose why PB concentration is positively corelated with nanofiber bundling and alignment, while also negatively correlated with bulk hydrogel diameter.

### Mechanical Characterization of Aligned Hydrogels

We next wanted to understand how differences in fibrillar nanostructure affect the mechanical properties of K2 hydrogels. We hypothesized that greater nanofibrillar alignment and packing would lead to stiffer K2 hydrogels. As K2 hydrogels are lifted from their gelation bath, hydrophilic cohesive forces counteract the hydrogel exiting the solution, and a downward tensile force is applied. We took advantage of this phenomenon to qualitatively compare the strength of K2 hydrogels fabricated into different gelation baths. Pre-gelled and 1X PB hydrogels cannot be removed from solution without fracturing (Movies S3 and S4), whereas 5X PB hydrogels (of large and small diameter) can be fully removed without fracture (Movies S5 and S6). These results were surprising, as only covalently crosslinked K2 hydrogels had previously been found to be strong enough to withstand be removed from solution.^32^ We therefore sought to further interrogate differences in mechanical properties between K2 gels.

In addition to being anisotropic and hydrated, the mechanical property regime of K2 hydrogels makes them challenging to characterize through most standard mechanical testing means. As a result, we employed optical coherence elastography (OCE),^33^ in which a 1 kHz non-contact microtap excitation was applied to one end of the cylindrical hydrogels, and the elastic wave speed was monitored as it propagated along the direction of nanofibrillar alignment (Figure 3a-d). Using this method, we could maintain K2 hydration and determine quantitative differences in stiffness between samples. The average wave speed through pre-gelled unaligned hydrogels was measured to be 0.80 m/s (Figure 3a and Movie S7), which was not significantly different from 1X PB hydrogels, which had an average wave speed of 0.71 m/s (Figure 3b and Movie S8). In contrast, the average wave speeds through 3X and 5X PB hydrogels were computed to be 1.9 (Figure 3c and Movie S9) and 5.5 m/s (Figure 3d and Movie S10), respectively. Together, these data show a positive correlation between PB concentration and wave speed along the direction of nanofibrillar alignment (Figure 3f). Assuming a constant density and using the Scholte wave model, the Young’s Moduli were calculated to be 2.0, 1.5, 11, and 99 kPa for the pre-gelled, 1X, 3X, and 5X PB hydrogels, respectively.

**Figure 3.**
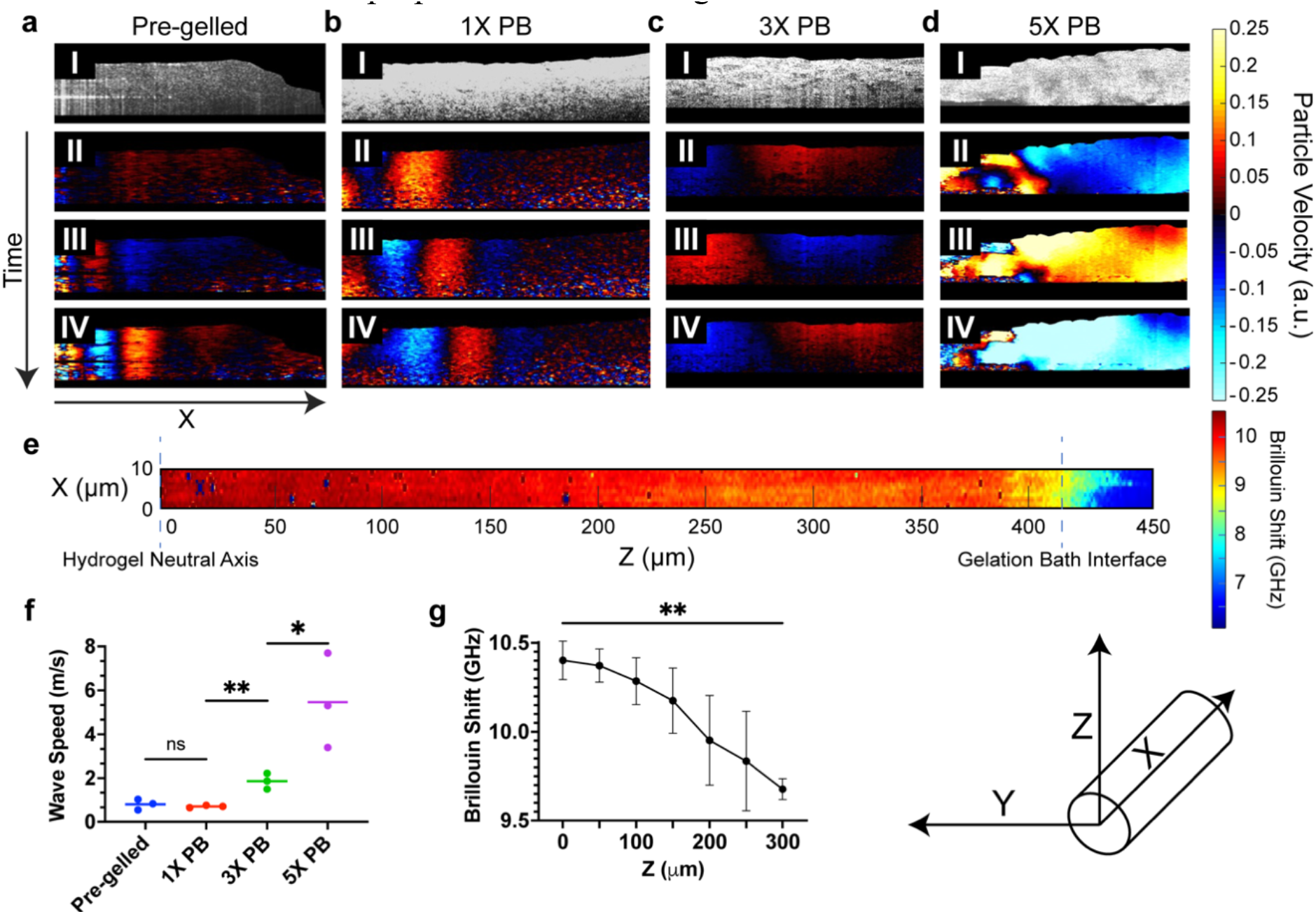
Mechanical Characterization of Aligned K2 Hydrogels. (a) Pre-gelled, (b) 1X PB, (c) 3X PB, and (d) 5X PB hydrogels (I) structural optical coherence tomography representations and (II) mechanical wave snapshots at t = 4.5 ms, (III) t = 5 ms, and (IV) 6.5 ms following a 1 kHz microtapping excitation. The color bar to the right indicates the particle velocity spectrum. (e) Brillouin shift profile for 5X PB hydrogel. The color bar to the right indicates the Brillouin shift (GHz). (f) Comparison of average wave speeds calculated using particle velocity profiles for each hydrogel type (n = 3 samples per condition; line at mean; *P < 0.05, **P < 0.01 by multiple Student’s *t* tests). (g) Comparison of Brillouin shift along the z-axis for 5X PB hydrogels (n = 3 samples with data pooled into 50 μm groupings; mean ± standard deviation; **P < 0.01 by Student’s *t* test).

We also imaged 5X PB hydrogels via Brillouin microscopy, which allows for an understanding of a material’s bulk modulus at a given location.^34^ We used this technique to understand how the hydrogel’s mechanical properties vary radially approaching the neutral axis (Figure 3e). The Brillouin shift was found to increase as a function of the Z-axis (Figure 3e,g and Figure S9), which suggests that the fiber alignment observed with SEM (Figure 2bIV) persists to the core. In addition, it agrees with the fact that shear forces during extrusion are highest along the neutral axis. Robust characterization of aligned K2 hydrogels using OCE and Brillouin microscopy allows us to conclude that higher nanofibrillar alignment leads to stiffer hierarchical matrices.

### Cellular Spreading on Aligned Hydrogels

We hypothesized that cells would recognize and align along the direction of K2 nanofibrillar alignment, and that higher scaffold alignment would better direct cellular alignment. Porcine valvular interstitial cells (VICs), which lie in aortic valve leaflets and are primarily responsible for secreting ECM, are known to respond to directional cues.^35–37^ We seeded them onto pre-gelled, 1X, 3X, and 5X PB hydrogels and observed high cellular viability for up to seven days (Figure S11). One day after seeding, confocal microscopy qualitatively revealed that VICs on 1X, 3X, and 5X PB hydrogels aligned along the direction of K2 nanofibrillar alignment, unlike the pre-gelled control (Figure 4a). Quantification of immunofluoresenctly labelled actin filaments (Figure 4dI) and cell nuclei (Figure 4dII) corroborated these observations (see methods for detailed descriptions). VICs maintained this alignment trend at day 3 (Figure 4b,e), and by day 7, VICs on 1X and 3X PB hydrogels directionally grew to confluency (Figure 4c,f). Further, spreading patterns of VICs on 1X PB hydrogels at day 3 (Figure 4g) and day 7 (Figure 4h) show that these scaffolds promote macroscopic cellular alignment. Surprisingly, VICs on 5X PB hydrogels aligned no better than those on the pre-gelled control at day 7 (Figure 4cIV,f). These results suggested that greater nanofibrillar alignment had no benefit, and further, that matrices with extremely high alignments were not able to direct cell spreading (Figure 1b).

**Figure 4.**
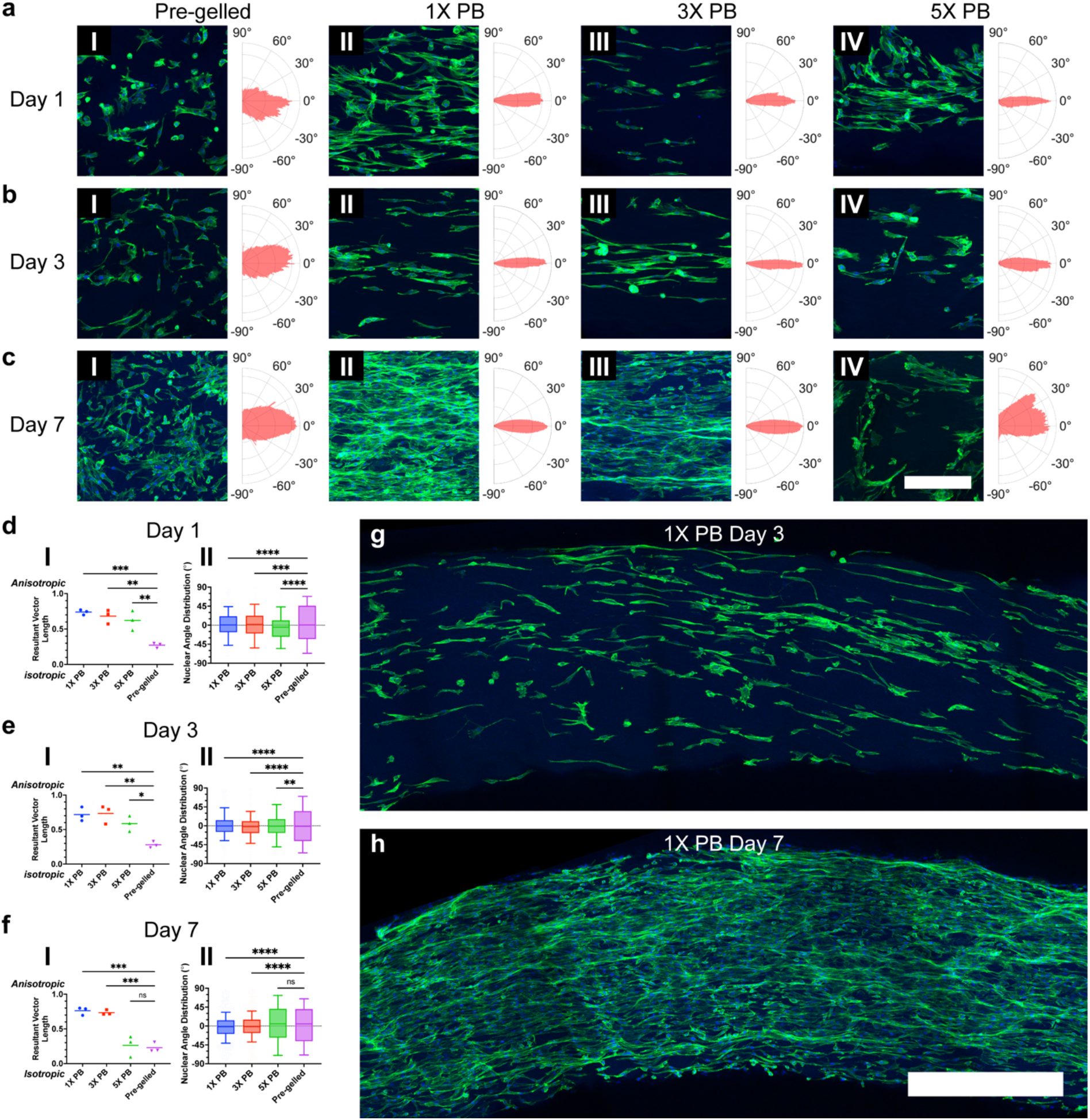
Valvular Interstitial Cell Spreading on Aligned K2 Hydrogels. (a) Day 1, (b) Day 3, and (c) Day 7 confocal microscopy images of VICs on (I) pre-gelled, (II) 1X PB, (III) 3X PB, and (IV) 5X PB hydrogels (DAPI = blue, F-actin = green; Scale bars = 300 μm). All images are maximum intensity projections of Z-stacks that have been cropped and rotated so the direction of K2 fibrous alignment is horizontal. Polar histograms to the right of each image display the Fourier gradient structure tensor calculated using the F-actin channels, where the direction of K2 fibrous alignment is 0°. (d) Day 1, (e) Day 3, and (f) Day 7 comparisons of (I) resultant vector lengths calculated from Fourier gradient structure tensors (n = 3 images; line at mean; *P < 0.05, **P < 0.01, ***P < 0.001 by one-way ANOVA and Dunnett’s multiple comparisons test) and (II) nuclear angle distributions with respect to the angle of K2 fibrous alignment (n = 3 images with between 294 and 1854 pooled nuclei; center-line, box bounds, and whiskers indicate the median, first and third quartiles, and 10 and 90 percentiles, respectively; **P < 0.01, ***P < 0.001, ****P < 0.0001 by Kolmogorov-Smirnov test). (g) Day 3 and (h) Day 7 confocal microscopy image stitches of VICs on 1X PB hydrogels (DAPI = blue, F-actin = green; Scale bar = 500 μm).

To corroborate these striking findings, we repeated this study using C2C12 cells, a murine myoblast cell line commonly used for skeletal muscle tissue engineering, which have also been shown to directionally align in response to mechanical cues.^10,38,39^ K2 hydrogels again supported high viability (Figure S12) and at day 1 after seeding, myoblasts aligned along the direction of K2 nanofibrillar alignment on 1X, 3X, and 5X PB hydrogels (Figure S13a,d). While C2C12 cells on 1X and 3X PB hydrogels maintained this alignment trend at day 3, cells on 5X PB hydrogels instead exhibited similar spreading patterns to those on pre-gelled unaligned controls (Figure S13b,e). By day 7, myoblasts on 1X and 3X PB hydrogels directionally grew to confluency (Figure S13c,f) and staining of myosin heavy chain at day 14 revealed that some myoblasts spontaneously differentiated into aligned myotubes on 1X and 3X PB hydrogels (Figure S14b,c). In contrast, myoblasts on 5X PB hydrogels spread orthogonal to the direction of K2 nanofibers (Figure S13cIV,fII) and aligned myotubes were not observed at day 14 (Figure S14d).

To further verify these results, we repeated VIC and C2C12 seeding studies onto K2 hydrogel scaffolds with half the diameter, supposing that hydrogel curvature may be affecting cellular growth. However, the cellular spreading patterns matched the previously observed trend (Figures S15 and S16). Finally, we examined whether hydrogel swelling could account for these results, but no differences in diameter were observed for 5X PB hydrogels when incubated in cell culture media for up to a week (Figure S17). Thus, cells on 1X and 3X PB hydrogels elongated in the direction of nanofibrillar alignment, but higher scaffold anisotropy led to no observable improvements in directing cells spreading, and the highly aligned 5X PB hydrogels did not promote cellular alignment (Figure 1b).

Subsequently, we imaged cellular spreading patterns via SEM to directly visualize how cells were interacting with the K2 matrices of varying alignments (Figure 5). Myoblasts and VICs were observed pulling on and becoming entangled in the 1X PB matrix (Figure 5a). In addition, both cell types were seen interacting with the 3X PB matrix similarly (Figure 5b), which was evident due to differences in matrix alignment directly next to, versus far from, cells (Figure 5bII). Cells on the 5X PB matrix had drastically different appearances and were not seen pulling on or disrupting the highly aligned matrix (Figure 5c). Instead, cells appeared to be spreading as if they were on 2D surface and astoundingly were even found to be spreading orthogonal to the direction of K2 nanofibrillar alignment (Figure 5cII).

**Figure 5.**
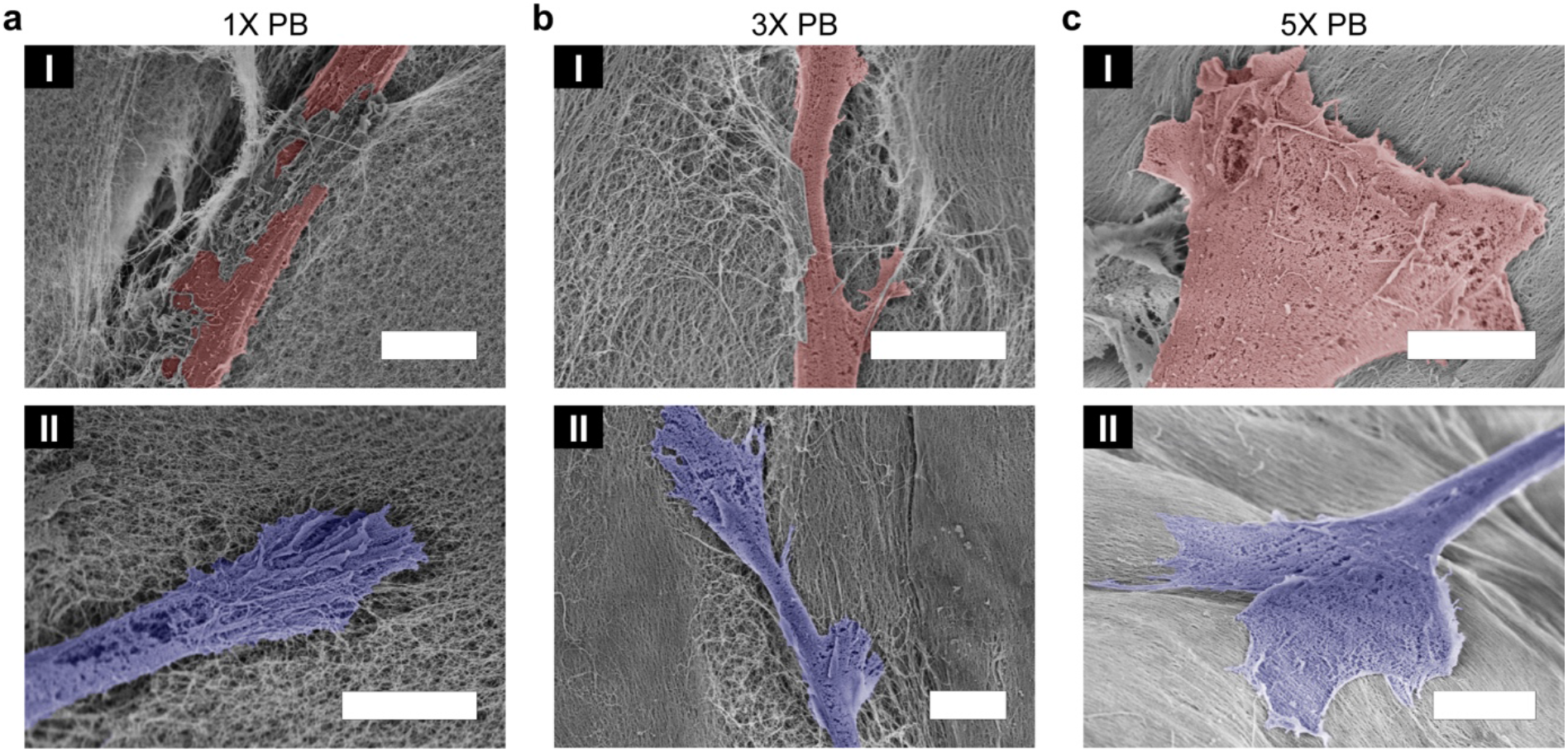
Cell-Matrix interactions on Aligned K2 Scaffolds. (a) 1X PB, (b) 3X PB, and (c) 5X PB hydrogel matrices with (I) C2C12 and (II) VICs observed via SEM (Scale bars = 5 μm). C2C12 and VIC cells are falsely colored red and blue, respectively.

After directly observing cell-matrix interactions, we can hypothesize **1**) why cells on 1X and 3X PB K2 hydrogels align similarly and **2**) why cells on 5X PB K2 hydrogels do not align along K2 alignment. Because K2 nanofibers are <10 nm in diameter (Figure S3d), we don’t observe cells interacting with individual fibers, but rather with the fibrous network. Cells are seen pulling on the 1X and 3X PB matrices similarly (Figure 5a,b), and differences in alignment and packing do not seem to impede this cellular behavior. We hypothesize that this pulling is a visual representation of mechanical coupling between the cell body and the underlying matrix and is what allows cells to sense and elongate along scaffold anisotropy. Thus for **1**, we believe that differences in alignment and packing between 1X and 3X PB matrices do not lead to significant differences in cells being able to mechanically couple with the K2 nanofibrous network, and therefore cells can sufficiently align on both matrices. In the case of **2**, we believe that K2 nanofibrous packing and bulk stiffness are too high for cells to effectively pull at and mechanically couple with the fibrous network, and therefore they cannot understand the mechanical alignment cues. Together, hypotheses for **1** and **2** allow us to speculate an explanation for the observed cellular spreading behaviors on K2 nanofibrous matrices and add more evidence towards the previously proposed “fibre recruitment” mechanism.^40^

## Conclusion

In conclusion, we have demonstrated an extrusion-based process to fabricate ECM-mimetic scaffolds of controlled alignments using self-assembling peptides. By coupling triggered gelation with an applied shear force, we uncovered a novel mechanism by which to tune nanofibrillar alignment within a supramolecular system. We rigorously characterized the structural and mechanical properties of the resulting hydrogels and demonstrated their ability to provide ECM-mimetic cues to guide directional cell growth. In addition, we were able to observe how nanoscale cell-matrix interactions dictate macroscopic cell spreading behaviors. This work has broad implications for the design of future self-assembling peptide systems and how cell-instructive fibrous cues can be used for bottom-up tissue engineering.

## Materials and Methods

### Peptide Synthesis and Purification

FMOC protected amino acids and Low loading rink amide MBHA resin were purchased from EMD Millipore (Burlington, MA). O-(7-Azabenzotriazol-1-yl)-N,N,N′,N′-tetramethyluronium hexafluorophosphate (HATU) was purchased from P3 BioSystems (Louisville, KY). N,N-dimethylformamide (DMF), dichloromethane (DCM), dimethyl sulfoxide (DMSO), piperidine, N,N-diisopropylethylamine (DiEA), acetic anhydride, trifluoroacetic acid (TFA), and diethyl ether were purchased from Fisher Scientific (Pittsburgh, PA). Piperidine, triisopropylsilane (TIPS), and anisole were purchased from Millipore Sigma (Burlington, MA).

K2 (full sequence: KKSLSLSLSLSLSLKK) was manually synthesized via solid phase peptide synthesis. Each coupling followed the same process: 2, 5-minute deprotections using 25% piperidine in DMF; 5, 30-second DMF washes; a ninhydrin test to confirm successful deprotection; a 20-minute coupling using 4 equivalents of an amino acid and 4 equivalents of HATU dissolved in 50% DMF/ 50% DMSO with 6 equivalents of DiEA; 2, 1-minute DCM washes; 2, 1-minute DMF washes; a ninhydrin test to confirm successful coupling. After coupling the last amino acid, one last deprotection was performed before acetylation of the N-terminus via 2, 45-minute couplings using an excess of DiEA and acetic anhydride in DCM. Following 3, 1-minute DCM washes, a ninhydrin test was used to confirm successful coupling. Peptide cleavage was performed for 3 hours using TFA with Milli-Q water, TIPS, and Anisole in excess as cleavage scavengers.

TFA was then evaporated off using Nitrogen gas, followed by trituration of the peptide in cold diethyl ether. Three cycles of centrifugation (10 minutes at 3400 RCF) followed by decanting of the supernatant were used to isolate the crude peptide. After allowing excess diethyl ether to evaporate overnight, the crude peptide was dissolved in Milli-Q water (0.5 – 1 wt%) and dialyzed against Milli-Q water for 4 days in 100–500 Dalton Spectra/Por Biotech Cellulose Ester Dialysis Membranes (Spectrum Laboratories Inc. Rancho Dominguez, CA). Next, the peptide solutions were pH adjusted to 7 and sterile filtered using 0.2 μm cellulose acetate sterile syringe filters (VWR International, Radnor, PA) under sterile conditions. Finally, peptides were frozen overnight at -80° C and lyophilized using a FreeZone 4.5 Liter Cascade Benchtop Freeze Dry System (Labconco Corporation, Kansas City, MO) for 3 days, before being transferred to a -20° C freezer for long term storage. Peptide synthesis was confirmed (Figure S1) using a Bruker AutoFlex Speed MALDI ToF (Bruker Instruments, Billerica, MA).

### Phosphate Buffer Gelation Baths

For each gelation bath, the masses of monosodium phosphate monohydrate and disodium phosphate heptahydrate were calculated to achieve the desired phosphate buffer concentration at pH 7. The buffer components and NaCl were then dissolved in Milli-Q water and sterile filtered using 0.2 μm cellulose acetate sterile syringe filters (VWR International, Radnor, PA) under sterile conditions.

### Polarized Light Microscopy (PLM)

PLM was performed on an Eclipse E400 (Nikon Corporation, Tokyo, Japan) equipped with cross polarizers. Images were captured on a mounted D7000 Digital Camera (Nikon Corporation, Tokyo, Japan) using a fixed exposure time and ISO. Samples were imaged on standard glass microscopy slides (Fisher Scientific, Pittsburgh, PA) at a consistent angle.

### Rheology

Rheology was performed using an AR-G2 rheometer (TA Instruments, New Castle, DE) equipped with a 12 mm parallel plate. 100 μL of 3% K2 in Milli-Q water was added to the stage and the plate was lowered to a gap of 500 μm. Excess solution was removed with a spatula and mineral oil was dripped around the stage to prevent dehydration of the sample during testing. Viscosity measurements were recorded during a shear sweep from 0.1 to 10 s^-1^.

### Cryo-Transmission Electron Microscopy (Cryo-TEM)

Cryo-TEM was performed using an FEI Tecnai F20 (FEI Company, Hillsboro, Oregon) equipped with a K2 summit camera (Gatan Inc, Pleasanton, CA). 3 wt% K2 in Milli-Q water was diluted in Milli-Q water to a final concentration of 0.1 wt% and added to glow discharged Quantifoil CUR 1.2/1.3 400 mesh grids (Electron Microscopy Sciences, Hatfield, PA). Samples were then frozen using a Vitrobot Mark IV Plunge System (Thermo Fisher Scientific, Waltham, MA) and loaded into a 626 Single tilt liquid nitrogen cryo-transfer holder (Gatan Inc, Pleasanton, CA). Cryo-transmission electron micrographs were captured at 200 kV. Brightness and contrast have been consistently modified to aid in visualization.

### Attenuated Total Reflectance Fourier Transform Infrared Spectroscopy (ATR-FTIR)

ATR-FTIR was performed using a Nicolet iS20 FT/IR spectrometer (Thermo Scientific, Waltham, MA) with a Golden Gate diamond window. 10 μL of 3 wt% K2 in Milli-Q water was added onto the window and dried using nitrogen. Spectra consisted of 30 accumulations at a resolution of 4 cm^−1^ with background subtraction. To compare relative heights in the amide I band at 1616 cm^−1^, spectra were normalized to the TFA peak at 1674 cm^−1^.

### Dynamic Light Scattering (DLS)

DLS was performed using a Malvern Zen 3600 Zetasizer (Malvern Instruments Ltd., Malvern, U.K.). 2 μL of 3 wt% K2 in Milli-Q water was diluted in Milli-Q water to a final concentration of 0.3 wt% and added to a disposable cuvette. Z-average and polydispersity index (PDI) were acquired using the default water parameters for all measurements.

### Scanning Electron Microscopy (SEM)

SEM was performed using a Helios NanoLab 660 Scanning Electron Microscope (FEI Company, Hillsboro, OR). K2 hydrogel samples were transferred to Porous Spec Pots (Electron Microscopy Sciences, Hatfield, PA) and subjected to a series of ethanol in Milli-Q water dilutions (30%, 50%, 60%, 70%, 80%, 90%, 2 × 100%), each for 10 minutes. Samples that also contained cells were fixed in 4% paraformaldehyde (Thermo Fisher Scientific, Waltham, MA) for 30 minutes prior to the serial dilution process. Next, samples were critical point dried using a Leica EM CPD300 (Leica Biosystems, Deer Park, IL) and coated with 5 nm of gold using a Denton Desk V Sputter System (Denton Vacuum, Moorestown, NJ). Scanning electron micrographs were captured at 1 - 2 kV, 25 pA and were cropped and rotated so the direction of K2 nanofibrillar alignment was horizontal.

### 2D Small-Angle X-Ray Scattering (2D SAXS)

2D SAXS was performed using a Rigaku S-MAX 3000 (Rigaku, Tokyo, Japan) at the University of Houston within the lab of Dr. Megan Robertson. K2 hydrogel samples were placed horizontally onto SpectroMembrane Polyimide 7.5 μm films (Chemplex Industries, Palm City, FL), mounted perpendicular to the x-ray path, and pulled to vacuum. After locating the X/Y location of maximum signal intensity, a 5-minute x-ray exposure was performed on each sample. Following background subtraction, the exposure pattern was integrated over the azimuthal angle using 1° increments, between the Q-range of 0.012 – 0.04 Å^-1^. Data points corresponding to the following angles were omitted from during plotting as they consistently presented large outliers: 1.5, 44.6, 88.7, 89.7, 90.8, 134.9, 152.9, 153.9, 177.9, 178.9, 180°.

### Swelling Measurements

After fabrication into their respective gelation baths using standard 100-1000 μL and 0.1-10 μL pipette tips (VWR International, Radnor, PA), K2 hydrogels were transferred to C2C12 growth media within a 12-well plate and stored in a 37° C incubator. Hydrogels were imaged using an OMAX 18 MP USB 3.0 Digital Camera (OMAX Microscope) at each time point and all diameter measurements were performed manually in ImageJ (National Institutes of health, Bethesda, MA).

### Optical Coherence Elastography (OCE)

Figure S10 shows a schematic of the custom-build OCE system. Wave excitation was generated using a 7x7x42 mm Piezo Stack Actuator (PiezoDrive, Newcastle, Australia) attached to a needle that was placed in direct contact with the hydrogel surface. Wave propagation was detected using a phase-sensitive Optical Coherence Tomography (PhS - OCT) system with ∼9 μm axial resolution (in air), ∼8 μm transversal resolution, and 0.28 nm of displacement stability. The A-line acquisition was set to 25 kHz during OCE acquisition. The piezo stack was driven by 5 pulses at 1 KHz, which was generated by a DG4162 Waveform Generator (RIGOL Technologies, Suzhou, China) and amplified by a PDu150 Three Channel 150 V piezo drive (PiezoDrive, Newcastle, Australia). The burst of the signal was synchronized with the OCT frame trigger during the M-B mode scan, to scan 500 points over 2.5 mm laterally with each M-mode scan consisting of 500 A-lines.

Axial particle velocities (*v*_!_) were calculated based on the depth dependent phase (φ) difference between two consecutive complex values A-lines, *v*_!_(*z, t*) = *Δ*φ(*z, t*)λ_0_/(4πΔ*t*), using *n* = 1.37 as the refractive index for phosphate buffered saline gelation baths,^41^ Δ*t* ≈ 40 *μs* (temporal resolution), and λ_0_ = 840 *nm* for the central wavelength of the OCT light source. Wave velocities were computed as the slope of the wave propagation on spatiotemporal images. Since K2 hydrogels were immersed in phosphate buffered saline gelation baths, the scan and wave propagation areas were under liquid. Therefore, the Scholte wave model was used to describe the elastic properties of the hydrogels where the shear wave velocity (*C*_s_) is related with the Scholte wave speed (*C*_sch_) by *C*_sch_ = 0.846*C*_s_.^42^ The Scholte Young’s modulus (*E*) was calculated with the formula: 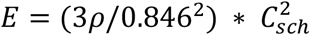, assuming a hydrogel density (ρ) of 1000 kg/m^3^. All calculations were performed in MATLAB 2020b (Mathworks Inc., Natick, MA).

### Brillouin Microscopy

The home-built Brillouin microscopy system was based on a two-stage virtually imaged phase array (VIPA) spectrometer. A single-mode 660 nm laser (Torus, Laser Quantum Inc., Fremont, CA) with an incident sample power of ∼17 mW was utilized, and a microscope was placed coaxially with the system to align the sample. A 40X water immersion microscope objective with a 0.8 numerical aperture was used to focus the laser beam onto the sample. The lateral resolution was ∼1.8 μm and the axial resolution was ∼2.3 μm. Before each experiment, calibration was performed using water, acetone, and methanol to acquire the spectral pixel resolution and the free spectral range. A 420 μm x 10 μm brillouin scan was taken with a step size of 1 μm. An electron-multiplying charged coupled device camera (iXon Andor, Belfast, U.K.) with an exposure time of 0.1 s was used to capture the backscattered light from the sample after passing through the VIPA-based spectrometer.

A program written in the LabVIEW 20.0.1 Development System (NI, Austin, TX) was used to synchronously control the CCD camera, 3D linear motorized stage, and data acquisition board. The obtained Brillouin spectrum was analysed using a Lorentzian fit to determine the Brillouin frequency shift. During the measurement, the sample was translated in 3D with the prescribed step size. Over a window, the Brillouin spectrum was summed before fitting to enhance the signal-to-noise ratio.^43^

### Cell Culture and Seeding

Fresh hearts from male and female young adult porcine (6 - 9 months old) were acquired from a commercial abattoir (Animal Technologies, Tyler, TX).^44^ The aortic valve cusps were dissected, soaked in 2.5% antibiotic/antimycotic solution (ABAM; stock concentration of 10,000 I.U. penicillin, 10,000 ug/mL streptomycin & 25 ug/mL amphotericin B; Corning, Corning, NY) in phosphate-buffered saline (PBS) and rinsed with PBS. The washing steps were repeated two times before the VICs were harvested following previously described methods.^36,45^ Briefly, the tissue was first incubated in an enzymatic solution containing 2 mg/mL collagenase II (Worthington Biochemical, Lakewood, NJ) in HyClone Low Glucose Dulbecco’s modified Eagle’s medium (Cytiva, Marlborough, MA) supplemented with 2.5% ABAM for 30 minutes under gentle agitation at 37° C. Next, the cusps were denuded of their endothelium before being minced and placed into an enzymatic mixture of 2 mg/mL collagenase III (Worthington Biochemical, Lakewood, NJ), 0.1 mg/mL hyaluronidase (Worthington Biochemical, Lakewood, NJ) and 2 mg/mL dispase II (STEMCELL Technologies, Vancouver, Canada) for 4 hours under gentle agitation at 37° C. The VICs were isolated from the digested tissue using a 70 μm cell strainer, pelleted at 1500 RCF for 5 minutes, resuspended and grown in 45% HyClone Low Glucose Dulbecco’s modified Eagle’s medium (Cytiva, Marlborough, MA) supplemented with 25 mM HEPES, 44% HyClone Ham’s Nutrient Mixture F12 (Cytiva, Marlborough, MA), 10% Gibco Fetal Bovine Serum (Thermo Fisher Scientific, Waltham, MA), and 1% penicillin-streptomycin (Thermo Fisher Scientific, Waltham, MA). Frozen C2C12 myoblasts were purchased from ATCC (Manassas, VA) and were grown in 89% Gibco Dulbecco’s modified Eagle’s medium (Thermo Fisher Scientific, Waltham, MA) supplemented with high glucose, sodium pyruvate, L-glutamine, and phenol red, 10% Gibco Fetal Bovine Serum (Thermo Fisher Scientific, Waltham, MA), and 1% penicillin-streptomycin (Thermo Fisher Scientific, Waltham, MA). All cells were used between passages 1 and 4.

For cell seeding studies, K2 hydrogels were fabricated under sterile conditions before being transferred to PBS. They were then moved to 8 Well high Bioinert μ-Slides (Ibidi, Gräfelfing, Germany) and placed into an incubator for 10 minutes prior to cell seeding. Cells were passaged to 200,000 cells/mL, PBS was removed from all wells to leave just the K2 hydrogels, and 10,000 cells (100,000 cells for cell viability studies) were added to the bottom of each well. Finally, media was added to each well and the slides were moved back into the incubator. Media was replaced every day for the remainder of the study.

### Staining + Imaging

Cells on K2 hydrogels were stained with Calcein AM and Ethidium homodimer (Thermo Fisher Scientific, Waltham, MA) following manufacturer protocols for cell viability studies. Imaging was performed using a Zeiss LSM800 Airyscan (Oberkochen, Germany) and the “3D objects counter” plugin within ImageJ (National Institutes of health, Bethesda, MA) was used to count live and dead cells, before being exported to Microsoft Excel (Microsoft Corporation, Redmond, WA) for viability calculations.

Immunostaining of cells on K2 hydrogels followed the process: 3 PBS washes; 30-minute fixation in 4 wt% paraformaldehyde (Thermo Fisher Scientific, Waltham, MA); 3 PBS washes; 10-minute quenching with 100 mM glycine; 30-minute permeabilization with 0.2% Triton-X (Fisher Scientific, Pittsburgh, PA) in PBS; 1-hour blocking with 1% Bovine Serum Albumin (Genetex, Irvine, CA) / 0.2% Triton-X in PBS; 1-hour staining with Alexa Fluor 488 Phalloidin (Thermo Fisher Scientific, Waltham, MA) diluted 1:400 in 1% BSA / 0.2% Triton-X in PBS; 3 PBS washes; 10-minute staining with DAPI (Thermo Fisher Scientific, Waltham, MA) diluted 1:500 in 1% BSA / 0.2% Triton-X in PBS; 3 PBS washes; clearing and storage in 88% Glycerol (Thermo Fisher Scientific, Waltham, MA). For immunostaining of C2C12 cell differentiation, Myosin Heavy Chain (MF 20, DSHB, Iowa City, IA) was diluted 1:10 and used as the primary antibody and Goat Anti-mouse Alexa Fluor Plus 488 (Thermo Fisher Scientific, Waltham, MA) was diluted 1:500 and used as the secondary antibody. All imaging was performed on a Zeiss LSM800 Airyscan (Oberkochen, Germany) and maximum intensity projections of z-stacks were created in ImageJ (National Institutes of health, Bethesda, MA). Collected images were cropped and rotated so the direction of K2 nanofibrillar alignment was always horizontal.

### Quantification of Actin and Nuclear Alignment

Unprocessed maximum intensity projections of immunofluoresenctly labelled actin filaments were imported into ImageJ (National Institutes of health, Bethesda, MA) and the plugin OrientationJ^46^ was used to locally determine the Fourier gradient structure tensor using a 2 pixel local window, a 5% minimum coherency, and a 5% minimum energy. The structure tensor was then imported into MATLAB 2023a (Mathworks Inc., Natick, MA) and the angle corresponding to the maximum value was shifted to be 0°. Next, a polar histogram was generated (which is displayed next to each confocal microscopy image) and the toolbox CircStat^47^ was used to calculate the mean resultant vector length of the polar distribution, where 0 corresponds to isotropy and 1 to anisotropy, as has been previously published.^48^

Unprocessed maximum intensity projections of immunofluoresenctly labelled cell nuclei were imported into ImageJ (National Institutes of health, Bethesda, MA) and the default automatic threshold was applied and converted to a mask. If the automated process failed, this step was performed manually. Next, the mask was dilated, eroded twice, and a watershed was applied. The Analyze Particle function was then used to find the angle of each ellipse and was exported into MATLAB 2023a (Mathworks Inc., Natick, MA). Finally, the difference between each ellipse angle and angle of K2 nanofibrillar alignment was calculated, similar to previously published methods.^13^

### Statistical Analysis

Prism 10 (GraphPad Software, Boston, MA) was used for all statistical analysis. Specifics about data plotted and statistical tests used are listed in all figure captions.

## Author contributions

A.C.F. and J.D.H. conceived the study. A.C.F. performed all PLM, rheology, ATR-FTIR, SEM, 2D SAXS, cell seeding studies, live/dead and immunostaining, confocal imaging, data processing including writing code and carrying out the quantification of actin and nuclear alignment, statistical analysis, and wrote the original manuscript draft. C.Z.D. performed OCE and completed data processing. L.T.Y. and C.C.C. prepared samples for Cryo-TEM and L.T.Y. performed all imaging. S.S. performed Brillouin microscopy and completed data processing. J.W.R.S. prepared samples for and performed DLS. J.T.M. created all gelation baths and measured sample diameter and swelling. A.J.T. synthesized and characterized K2. E.G.H. isolated VIC cells. K.J.G.A. oversaw the work of E.G.H. M.S. and K.V.L. oversaw the work of C.Z.D. and S.S. J.D.H. oversaw the work of A.C.F., L.T.Y., J.W.R.S., J.T.M., A.J.T., and C.C.C. All authors reviewed and edited the manuscript.

## Notes

A.C.F. and J.D.H. are listed as co-inventors on pending U.S. patent application #63/510,818. K.V.L. and M.S. have a financial interest in ElastEye LLC, which is not directly related to this work. The remaining coauthors declare they have no competing interests.

## Acknowledgments

This work was supported by National Institutes of Health grants R01DE021798 (to J.D.H.), R01EY022362 (to K.V.L.), R01HD095520 (to K.V.L.), R01EY030063 (to K.V.L.); National Science Foundation grant 2129122 (to K.J.G.A.); National Science Foundation Graduate Research Fellowship Program (to A.C.F., J.W.R.S., C.C.C.). We thank M. Robertson for access to the Rigaku S-MAX 3000 2D-SAXS at the University of Houston and J. Hanson for training. We also thank B. Utama for help with confocal microscopy and Rice University’s Biomaterials Lab. We thank J. Miller and J. Beckham for helpful discussions.

