## Supporting Information for "Tunable Macroscopic Alignment of Self-Assembling Peptide Nanofibers"

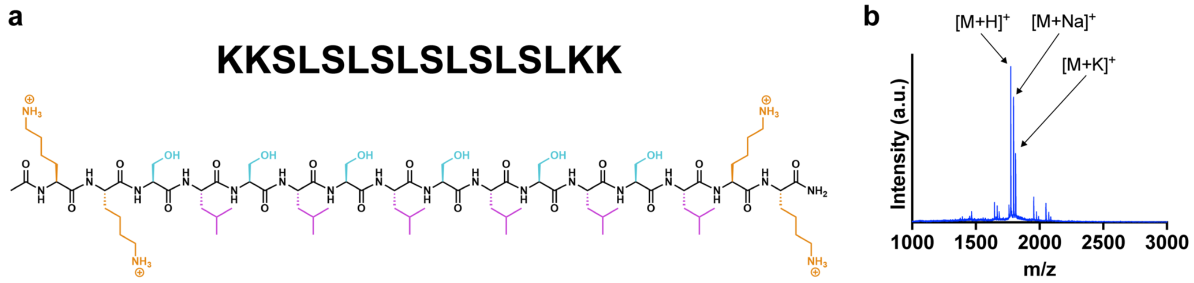


**Figure S1.** K2 Characterization. (a) K2 sequence and structure. (b) Matrix-assisted laser desorption/ionization (MALDI) spectrum showing successful synthesis.

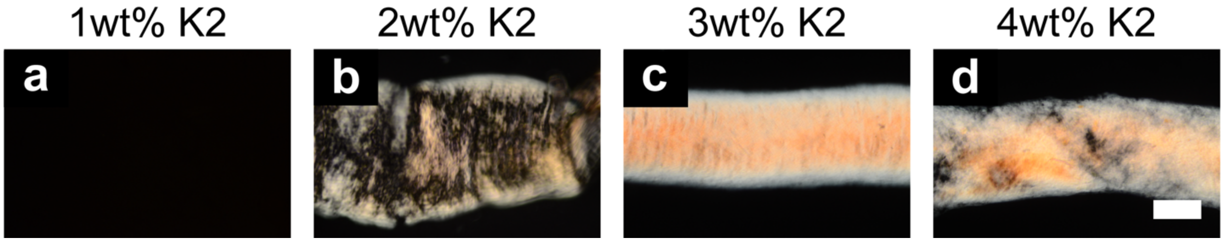


**Figure S2.** K2 Concentration Screen. Polarized light microscopy of (a) 1wt%, (b) 2wt%, (c) 3wt%, and (d) 4wt% K2 in Milli-Q water precursor solutions extruded through a 100-1000 μL pipette tip into a phosphate-buffered saline gelation bath (scale bar = 500 μm).

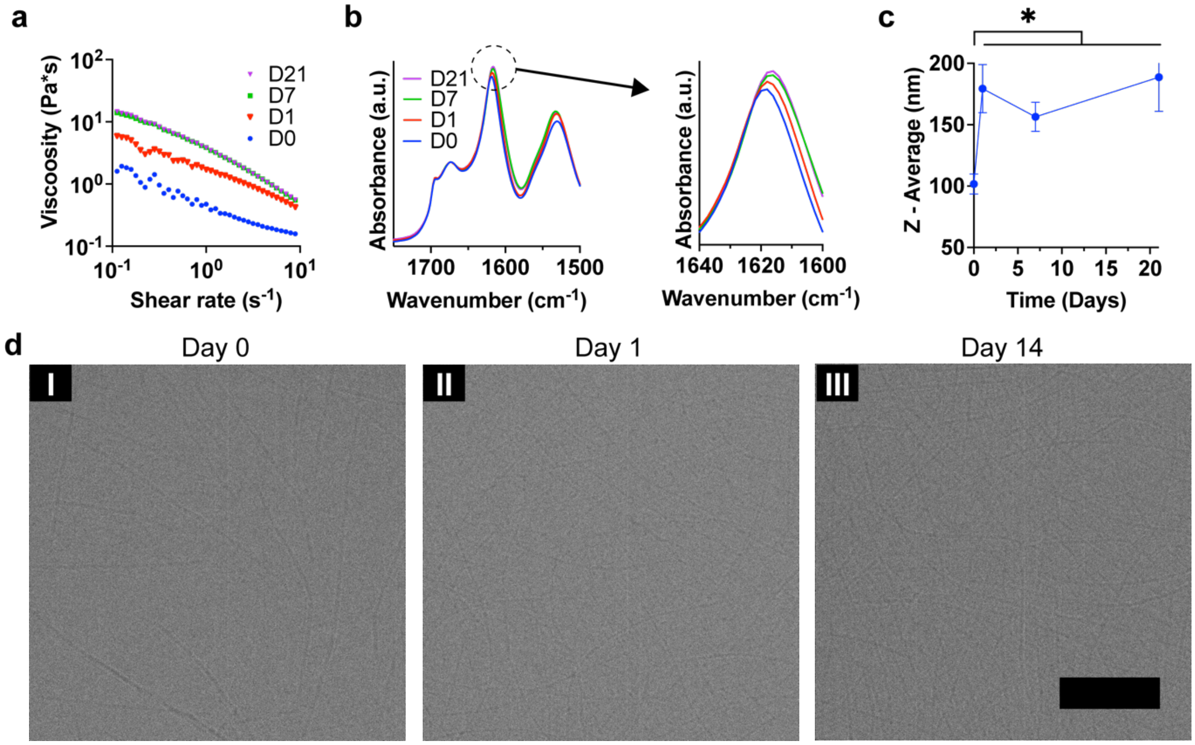


**Figure S3.** Precursor Solution Equilibration. (a) Rheology shear sweep between 0.1 and 10 s-1, (b) Attenuated total reflectance Fourier transform infrared spectroscopy between 1500 and 1750 cm-1, and (c) dynamic light scattering z-average measurements (n = 3; mean ± standard deviation; *P < 0.05 by one-way ANOVA and Tukey's multiple comparisons test) of 3wt% K2 in Milli-Q water precursor solution at days 0, 1, 7, and 21. (d) Cryo-transmission electron microscopy of 3wt% K2 in Milli-Q water precursor solution at days (I) 0, (II) 1, and (III) 14 (scale bar = 200 nm).

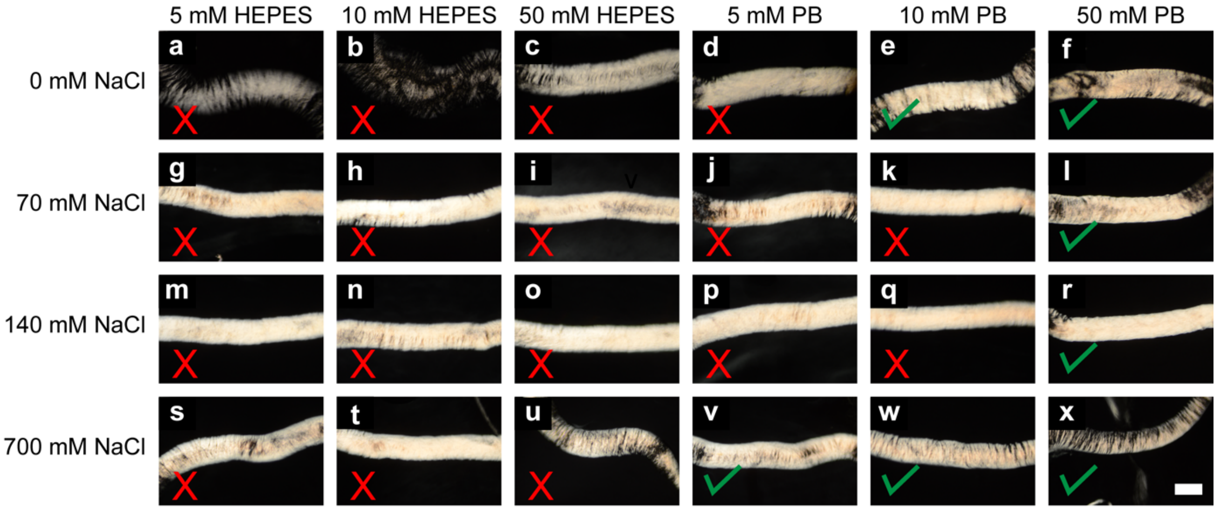


**Figure S4.** Gelation Bath Composition Screen. Polarized light microscopy of 3wt% K2 in Milli-Q water precursor solution extruded through a 0.1-10 µL pipette tip into a series of pH 7 gelation baths (scale bar = 500 µm). Green checkmarks indicate that hydrogels could be lifted from solution without fracturing, whereas red X's indicate that hydrogels fractured when lifted from solution. (a) 5 mM HEPES/ 0 mM NaCl. (b) 10 mM HEPES/ 0 mM NaCl. (c) 50 mM HEPES/ 0 mM NaCl. (d) 5 mM phosphate buffer (PB)/ 0 mM NaCl. (e) 10 mM PB/ 0 mM NaCl. (f) 50 mM PB/ 0 mM NaCl. (g) 5 mM HEPES/ 70 mM NaCl. (h) 10 mM HEPES/ 70 mM NaCl. (i) 50 mM HEPES/ 70 mM NaCl. (j) 5 mM PB/ 70 mM NaCl. (k) 10 mM PB/ 70 mM NaCl. (l) 50 mM PB/ 70 mM NaCl. (m) 5 mM HEPES/ 140 mM NaCl. (n) 10 mM HEPES/ 140 mM NaCl. (o) 50 mM HEPES/ 140 mM NaCl. (p) 5 mM PB/ 140 mM NaCl. (q) 10 mM PB/ 140 mM NaCl. (r) 50 mM PB/ 140 mM NaCl. (s) 5 mM HEPES/ 700 mM NaCl. (t) 10 mM HEPES/ 700 mM NaCl. (u) 50 mM HEPES/ 700 mM NaCl. (v) 5 mM PB/ 700 mM NaCl. (w) 10 mM PB/ 700 mM NaCl. (x) 50 mM PB/ 700 mM NaCl.

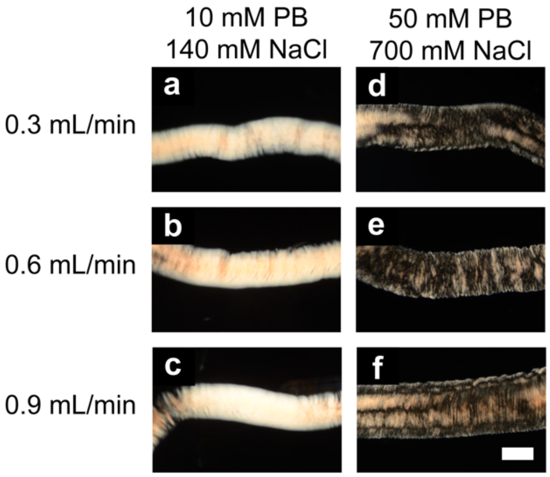


**Figure S5.** Precursor Solution Extrusion Rate Screen. Polarized light microscopy of 3wt% K2 in Milli-Q water precursor solution extruded at (a, d) 0.3 mL/min, (b, e) 0.6 mL/min, and (c, f) 0.9 mL/min into (a-c) 10 mM phosphate buffer (PB)/ 140 mM NaCl, pH 7 and (d-f) 50 PB/ 700 mM NaCl, pH 7 gelation baths (scale bar = 500 µm).

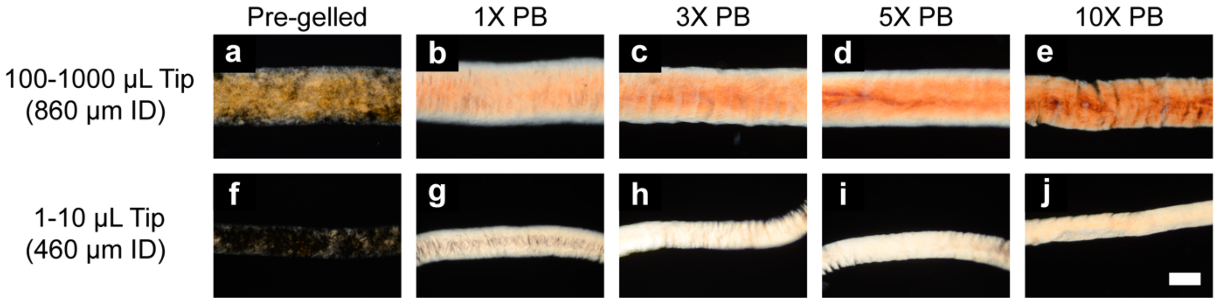


**Figure S6.** Gelation Bath Phosphate Buffer Screen. Polarized light microscopy of (a, f) pre-gelled, (b, g) 1X PB, (c, h) 3X PB, (d, i) 5X PB, and (e, j) 10X PB hydrogels (scale bar = 500 µm). (a-e) 100-1000 µL and (f-j) 0.1-10 µL pipette tips were used to manually extrude the precursor solutions.

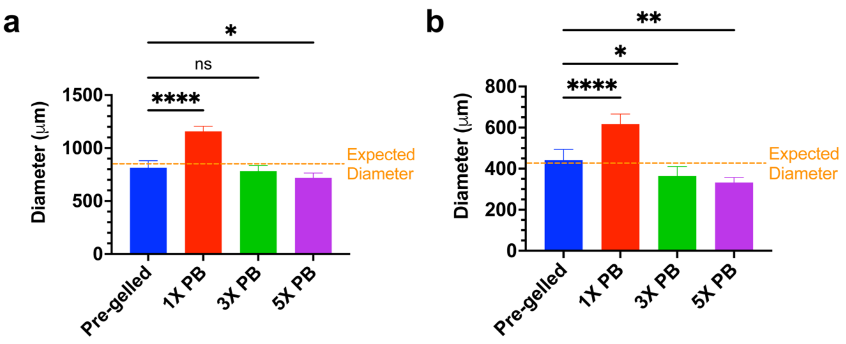


**Figure S7.** Bulk K2 Hydrogel Geometry. Hydrogel diameter measurements of K2 hydrogels fabricated through (a) 100-1000 μL (n = 6 samples; mean ± standard deviation; $*P < 0.05$, $****P < 0.0001$ by one-way ANOVA and Dunnett's multiple comparisons test) and (b) 0.1-10 μL pipette tips (n = 6 samples; mean ± standard deviation; $*P < 0.05$, $**P < 0.01$, $****P < 0.0001$ by one-way ANOVA and Dunnett's multiple comparisons test). The expected diameters for the large and small hydrogels are indicated in orange and are 860 and 430 μm, respectively.

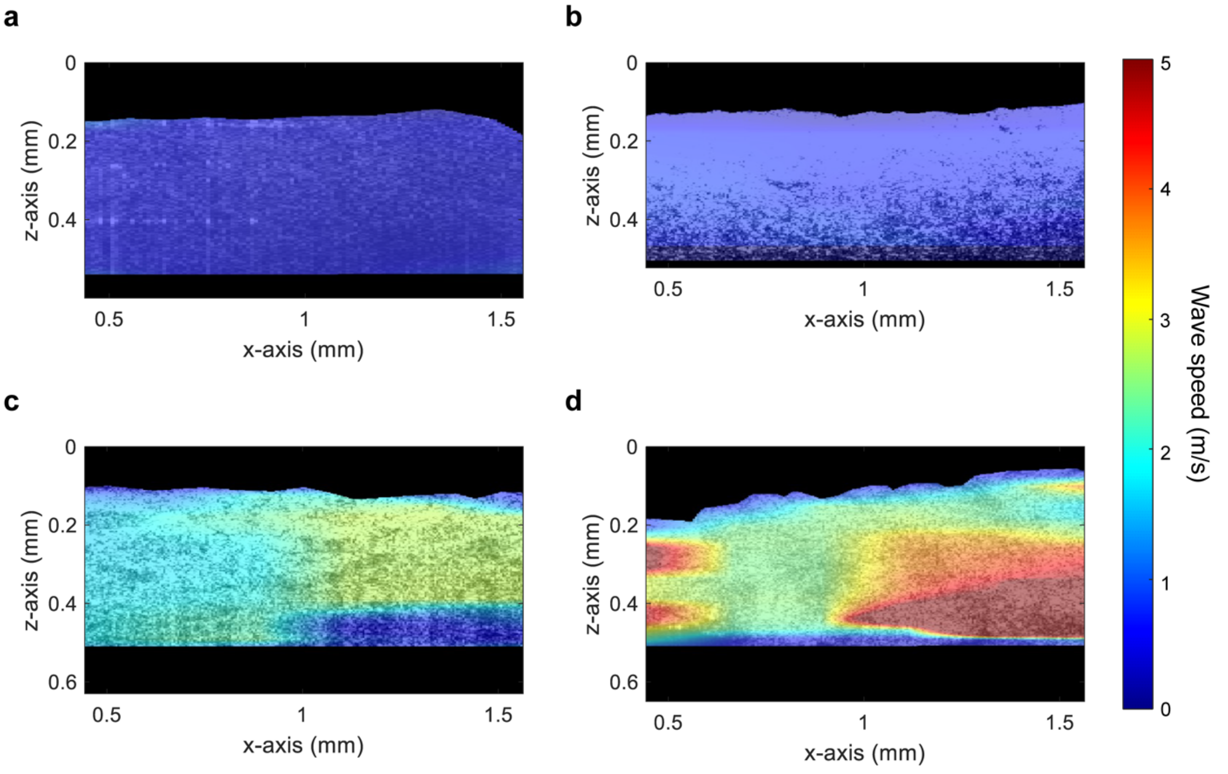


**Figure S8.** Stiffness of Aligned K2 Hydrogels. Optical Coherence Elastography elastic wave speed maps for (a) pre-gelled, (b) 1X PB, (c) 3X PB, and (d) 5X PB hydrogels. The color bar to the right indicates wave speed (m/s).

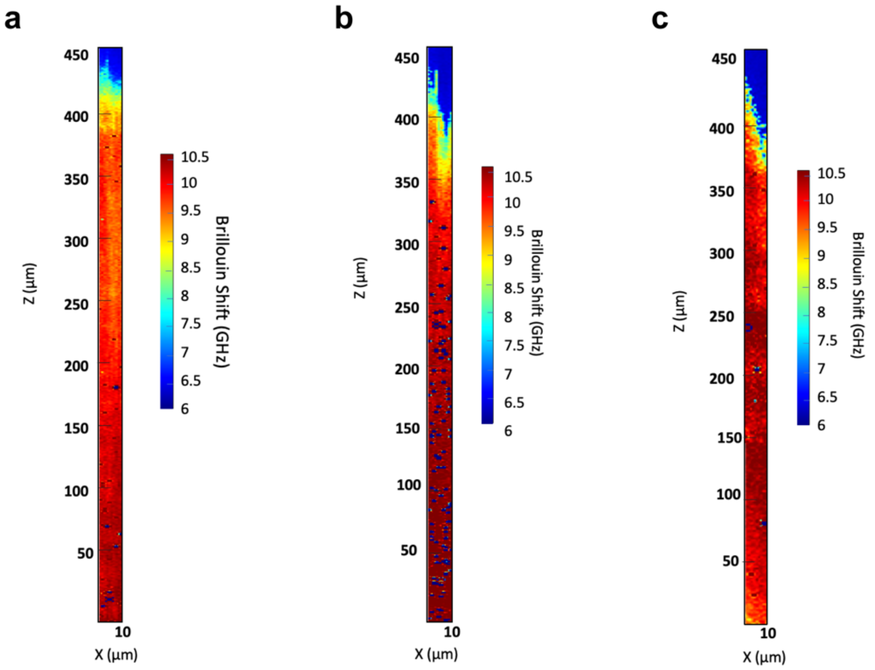


**Figure S9.** Brillouin Shift Profiles for 5X PB Hydrogels. (a-c) Brillouin frequency shift maps of three 5X PB hydrogels. The color bar to the right indicates the Brillouin shift (GHz).

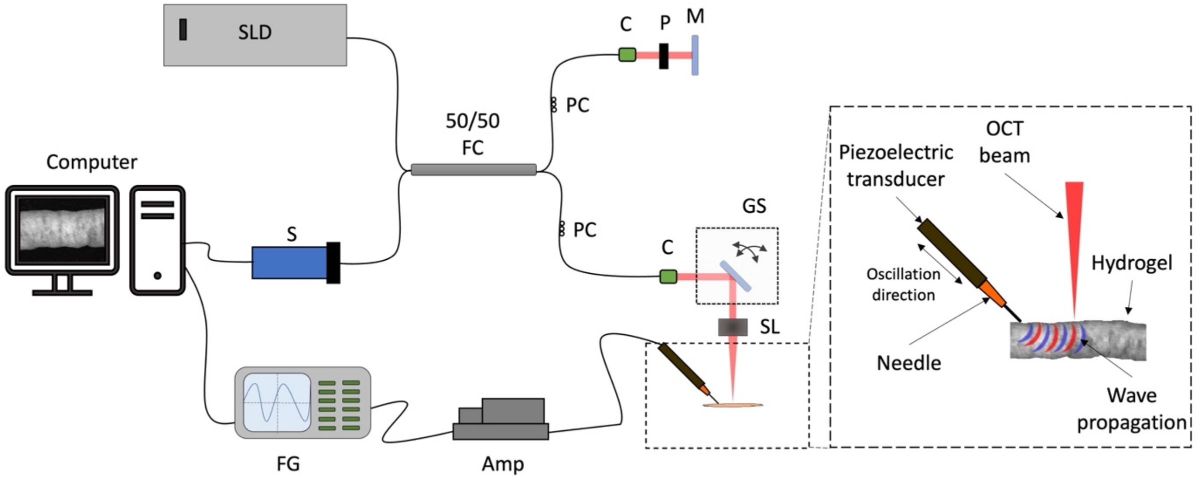


**Figure S10.** Optical Coherence Elastography System Schematic. Right inset shows the hydrogel alignment scanned by the OCT beam and the piezoelectric transducer with an attached needle producing a wave propagation parallel to the OCT scanning direction. 2D OCE data was acquired by the 2D galvo scanner motion in the x-z plane. C, collimator; FC, fiber coupler; GS, 2D galvo scanner; P, pinhole; PC, polarization controller; S, spectrometer; Amp, RF amplifier; and SLD, a superluminescent diode.

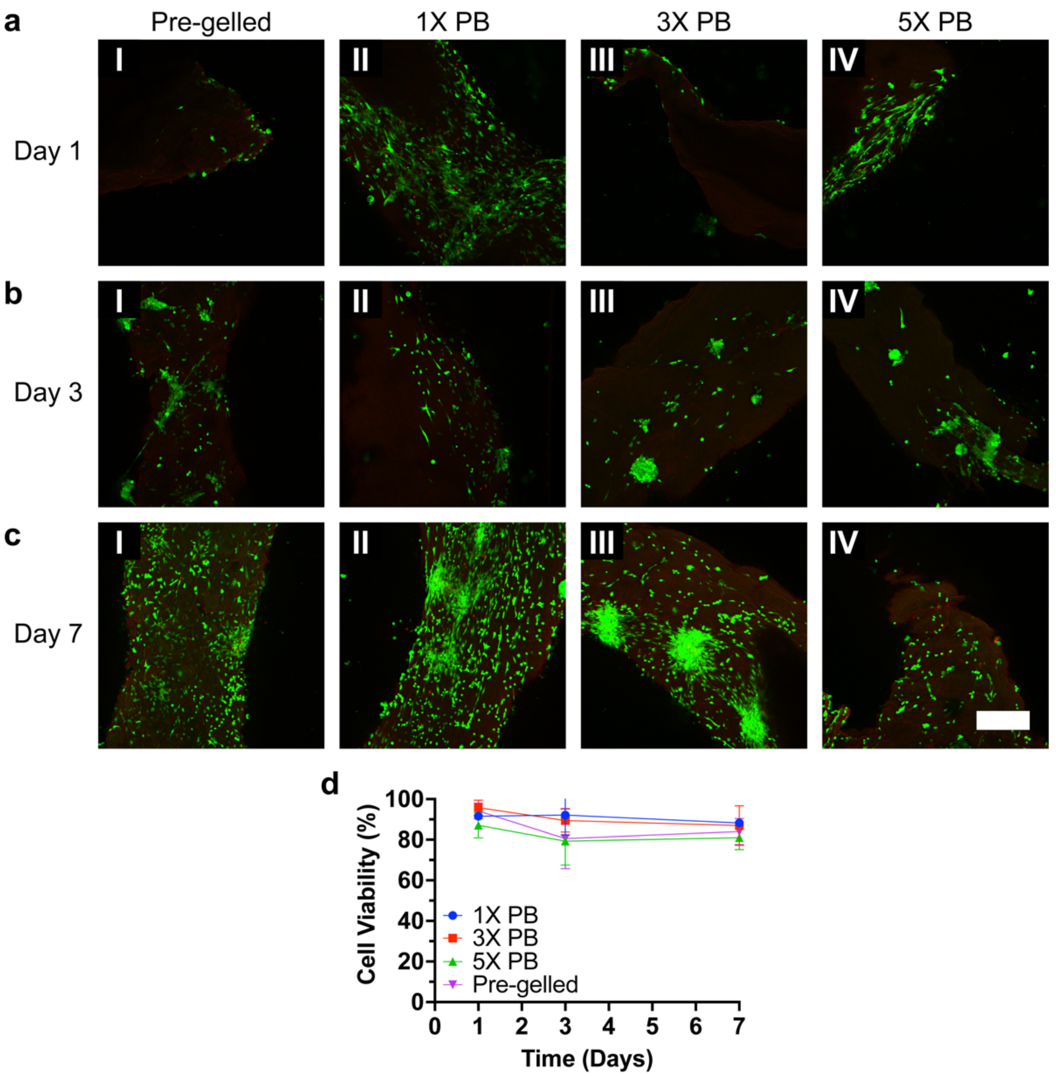


**Figure S11.** VIC Cell Viability on K2 Hydrogels. (a) Day 1, (b) Day 3, and (c) Day 7 confocal microscopy images of VICs on (I) pre-gelled, (II) 1X PB, (III) 3X PB, and (IV) 5X PB hydrogels (Calcein AM = green, Ethidium homodimer = red; scale bar = 300 µm). All images are maximum intensity projections of Z-stacks. (d) Cell viability calculations derived from confocal microscopy images (n = 3 images per time point per group; mean ± standard deviation).

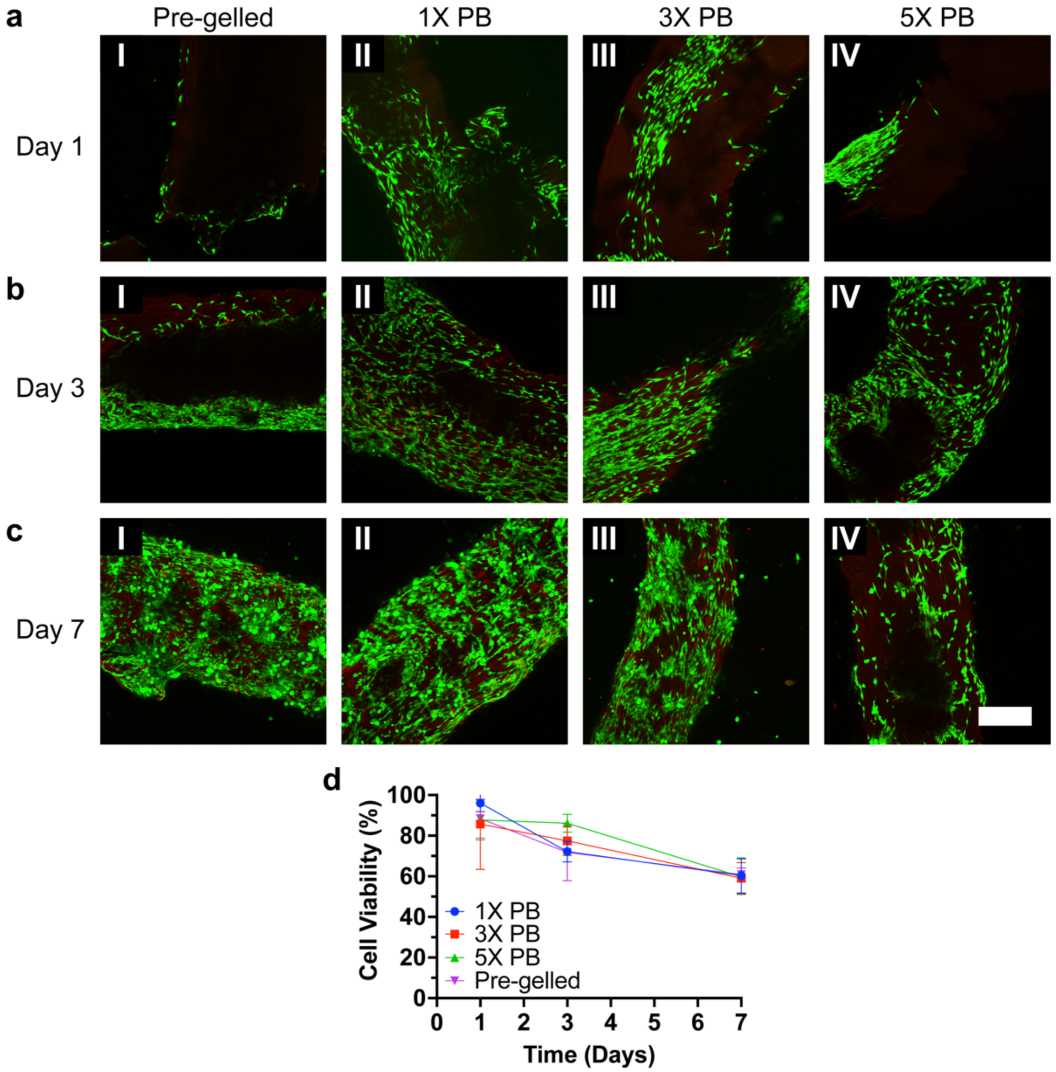


**Figure S12.** C2C12 Cell Viability on K2 Hydrogels. (a) Day 1, (b) Day 3, and (c) Day 7 confocal microscopy images of C2C12 cells on (I) pre-gelled, (II) 1X PB, (III) 3X PB, and (IV) 5X PB hydrogels (Calcein AM = green, Ethidium homodimer = red; scale bar = 300 μm). All images are maximum intensity projections of Z-stacks. (d) Cell viability calculations derived from confocal microscopy images (n = 3 images per time point per group; mean ± standard deviation).

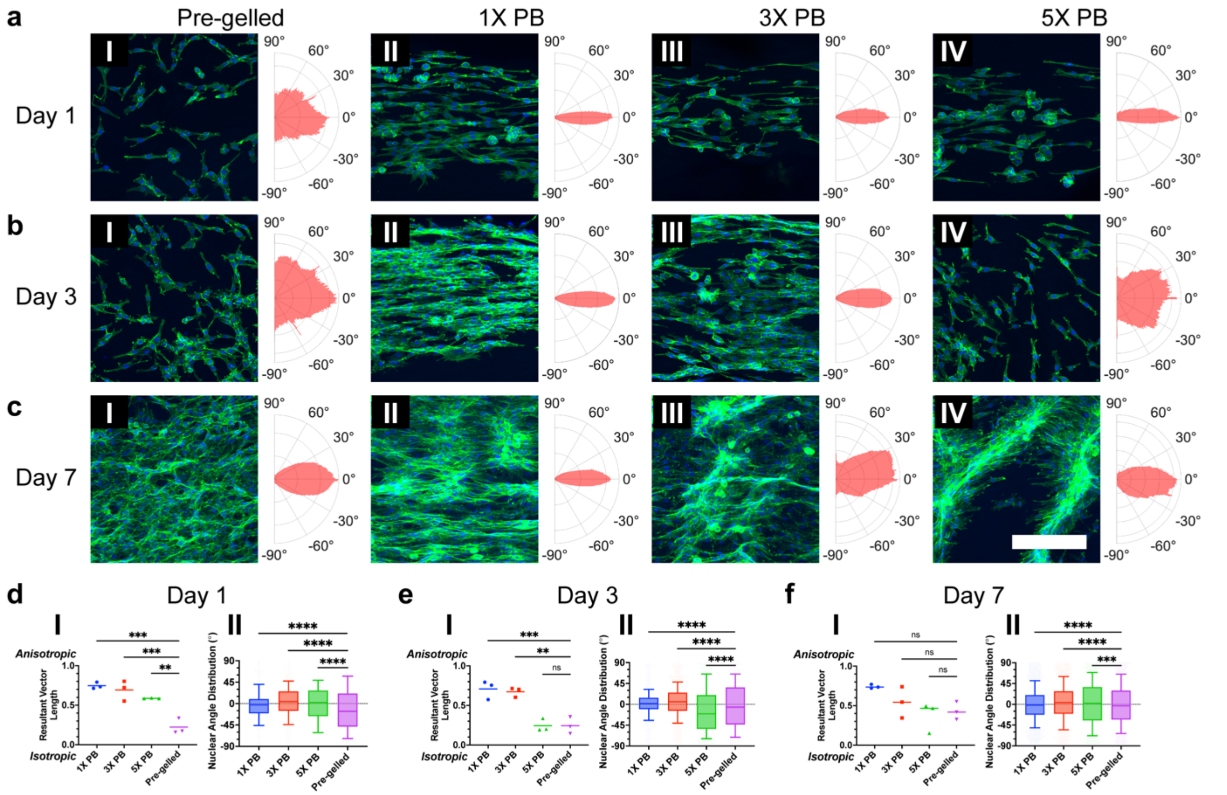

**Figure S13.** C2C12 Cell Spreading on K2 Hydrogels. (a) Day 1, (b) Day 3, and (c) Day 7 confocal microscopy images of C2C12 cells on (I) pre-gelled, (II) 1X PB, (III) 3X PB, and (IV) 5X PB hydrogels (DAPI = blue, F-actin = green; scale bar = 300 µm). All images are maximum intensity projections of Z-stacks that have been cropped and rotated so the direction of K2 fibrous alignment is horizontal. Polar histograms to the right of each image display the Fourier gradient structure tensor calculated using the F-actin channels, where the direction of K2 fibrous alignment is 0°. (d) Day 1, (e) Day 3, and (f) Day 7 comparisons of (I) resultant vectors lengths calculated from Fourier gradient structure tensors (n = 3 images; line at mean; $^{**}P < 0.01$, $^{***}P < 0.001$ by one-way ANOVA and Dunnett's multiple comparisons test) and (II) nuclear angle distributions with respect to the angle of K2 fibrous alignment (n = 3 images with between 242 and 2827 pooled nuclei; center-line, box bounds, and whiskers indicate the median, first and third quartiles, and 10 and 90 percentiles, respectively; $^{***}P < 0.001$, $^{****}P < 0.0001$ by Kolmogorov-Smirnov test).

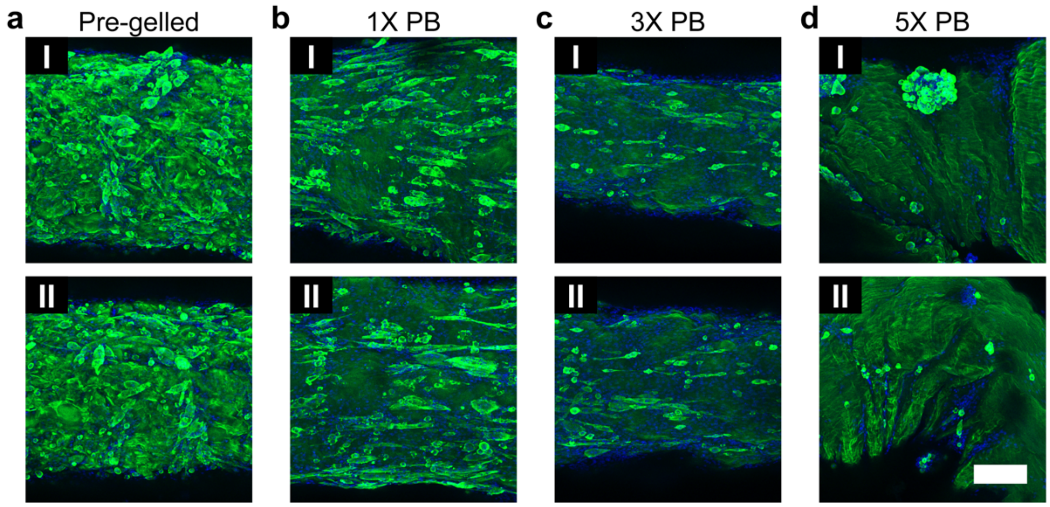


**Figure S14.** C2C12 Cell Differentiation on K2 Hydrogels. Confocal microscopy images of C2C12 cells on (a) pre-gelled, (b) 1X PB, (c) 3X PB, and (d) 5X PB hydrogels (DAPI = blue, Myosin heavy chain = green; scale bar = 300 μm). All images are maximum intensity projections of Z-stacks that have been cropped and rotated so the direction of K2 fibrous alignment is horizontal.

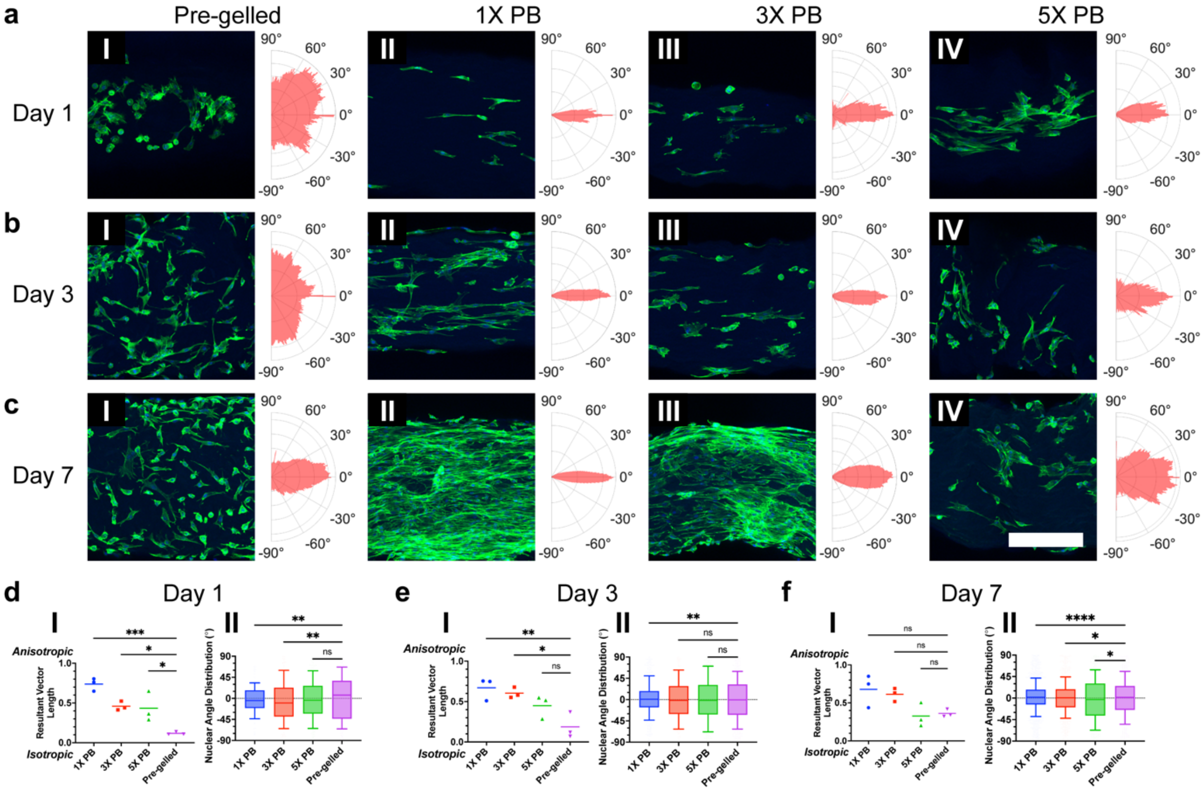


**Figure S15.** VIC Cell Spreading on Small Diameter K2 Hydrogels. (a) Day 1, (b) Day 3, and (c) Day 7 confocal microscopy images of VICs on (I) pre-gelled, (II) 1X PB, (III) 3X PB, and (IV) 5X PB hydrogels (DAPI = blue, F-actin = green; scale bar = 300 µm) fabricated using 0.1-10 µL pipette tips. All images are maximum intensity projections of Z-stacks that have been cropped and rotated so the direction of K2 fibrous alignment is horizontal. Polar histograms to the right of each image display the Fourier gradient structure tensor calculated using the F-actin channels, where the direction of K2 fibrous alignment is 0°. (d) Day 1, (e) Day 3, and (f) Day 7 comparisons of (I) resultant vectors lengths calculated from Fourier gradient structure tensors (n = 3 images; line at mean; *$P < 0.05$, **$P < 0.01$, ***$P < 0.001$ by one-way ANOVA and Dunnett's multiple comparisons test) and (II) nuclear angle distributions with respect to the angle of K2 fibrous alignment (n = 3 images with between 113 and 1416 pooled nuclei; center-line, box bounds, and whiskers indicate the median, first and third quartiles, and 10 and 90 percentiles, respectively; *$P < 0.05$, **$P < 0.01$, ****$P < 0.0001$ by Kolmogorov-Smirnov test).

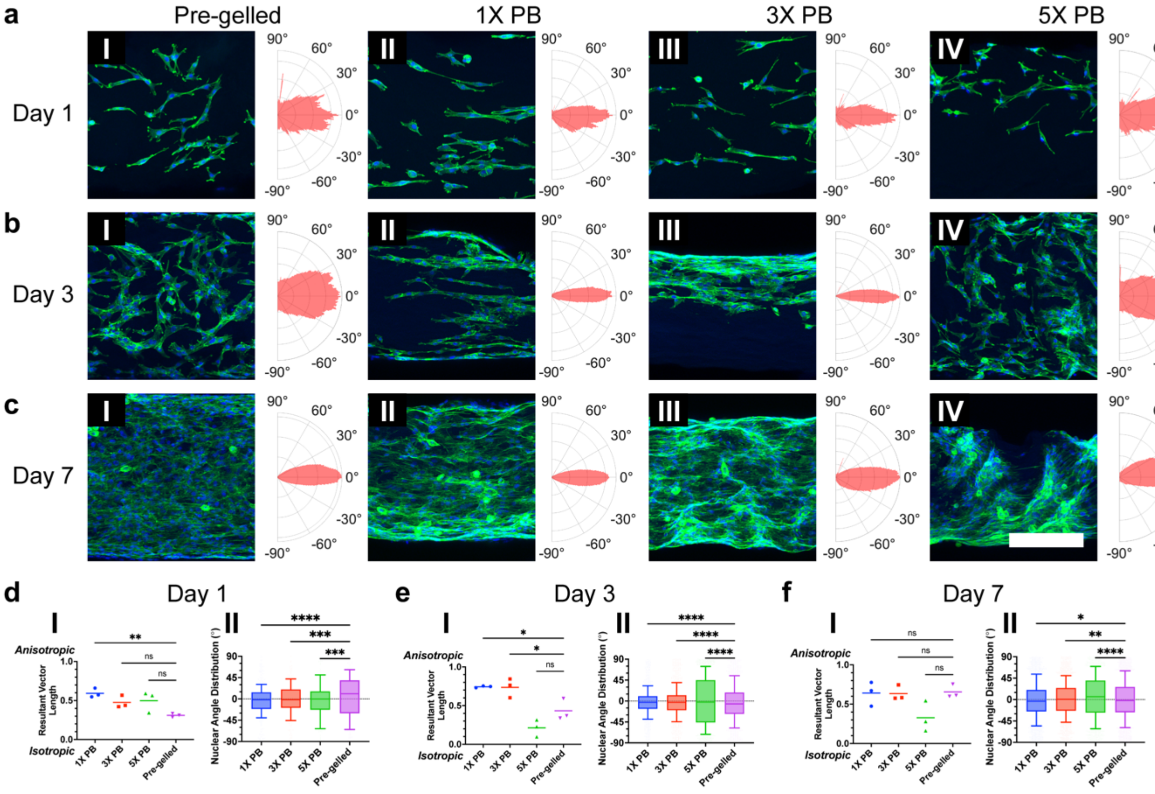


**Figure S16.** C2C12 Cell Spreading on Small Diameter K2 Hydrogels. (a) Day 1, (b) Day 3, and (c) Day 7 confocal microscopy images of C2C12 cells on (I) pre-gelled, (II) 1X PB, (III) 3X PB, and (IV) 5X PB hydrogels (DAPI = blue, F-actin = green; scale bar = 300 μm) fabricated using 0.1-10 μL pipette tips. All images are maximum intensity projections of Z-stacks that have been cropped and rotated so the direction of K2 fibrous alignment is horizontal. Polar histograms to the right of each image display the Fourier gradient structure tensor calculated using the F-actin channels, where the direction of K2 fibrous alignment is 0°. (d) Day 1, (e) Day 3, and (f) Day 7 comparisons of (I) resultant vectors lengths calculated from Fourier gradient structure tensors (n = 3 images; line at mean; $*P < 0.05$, $**P < 0.01$ by one-way ANOVA and Dunnett's multiple comparisons test) and (II) nuclear angle distributions with respect to the angle of K2 fibrous alignment (n = 3 images with between 128 and 2437 pooled nuclei; center-line, box bounds, and whiskers indicate the median, first and third quartiles, and 10 and 90 percentiles, respectively; $*P < 0.05$, $**P < 0.01$, $***P < 0.001$, $****P < 0.0001$ by Kolmogorov-Smirnov test).

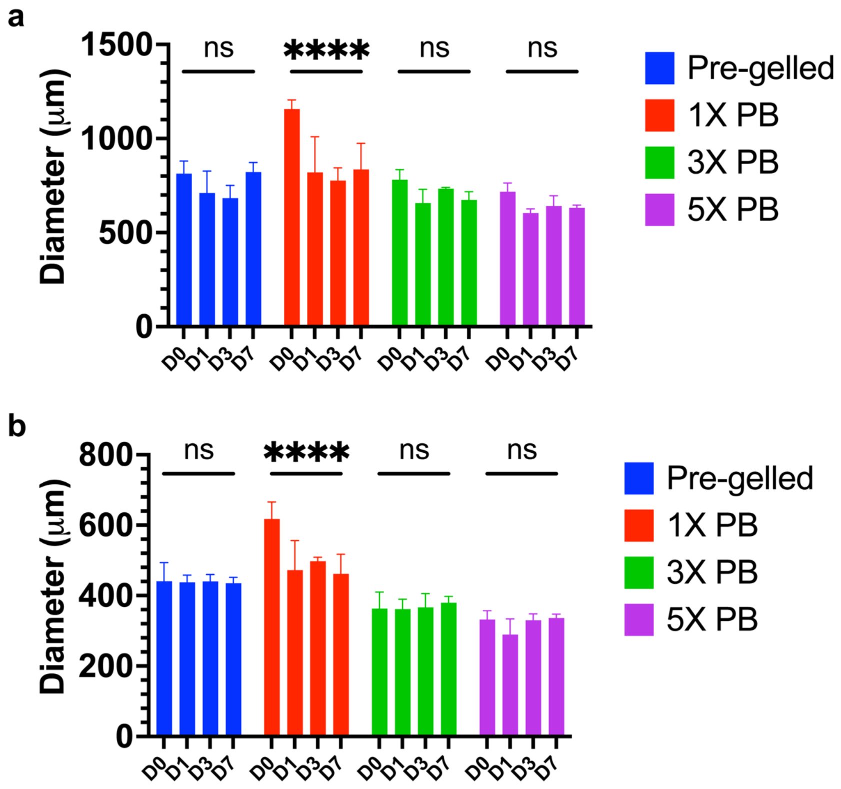


**Figure S17.** K2 Hydrogel Swelling. Hydrogel diameter measurements of K2 hydrogels fabricated through (a) 100-1000 µL (n = 2 - 6 samples per time point; mean ± standard deviation; ****P < 0.0001 by two-way ANOVA and Dunnett's multiple comparisons test) and (b) 0.1-10 µL pipette tips (n = 3 - 6 samples per time point; mean ± standard deviation; ****P < 0.0001 by two-way ANOVA and Dunnett's multiple comparisons test). The expected diameters for the large and small hydrogels are 860 and 430 µm, respectively.

Movie S1.
Hydrogel Fabrication

Movie S2.
Hydrogel Fabrication PLM

Movie S3.
Pre-gelled K2 Hydrogel Pick Up

Movie S4.
1X PB K2 Hydrogel Pick Up

Movie S5.
5X PB K2 Hydrogel Large Pick Up

Movie S6.
5X PB K2 Hydrogel Small Pick Up

Movie S7.
Pre-gelled K2 Hydrogel OCE

Movie S8.
1X PB K2 Hydrogel OCE

Movie S9.
3X PB K2 Hydrogel OCE

Movie S10.
5X PB K2 Hydrogel OCE
